# Neural Extracellular Matrix Remodeling Signatures in Genetic and Acquired Mouse Models of Epilepsy

**DOI:** 10.1101/2023.04.19.537468

**Authors:** Armand Blondiaux, Shaobo Jia, Anil Annamneedi, Gürsel Çalışkan, Jana Schulze, Carolina Montenegro-Venegas, Robert C. Wykes, Anna Fejtova, Matthew C. Walker, Oliver Stork, Eckart D. Gundelfinger, Alexander Dityatev, Constanze I. Seidenbecher

## Abstract

Epilepsies are multifaceted neurological disorders characterized by abnormal brain activity, e.g., caused by imbalanced synaptic excitation and inhibition. The neural extracellular matrix (ECM) is dynamically modulated by physiological and pathophysiological activity and critically involved in controlling the brain’s excitability. We used different epilepsy models, i.e. mice lacking the presynaptic scaffolding protein Bassoon at excitatory, inhibitory or all synapse types as genetic models for rapidly generalizing early-onset epilepsy, and intra-hippocampal kainate injection, a model for acquired temporal lobe epilepsy, to study the relationship between epileptic seizures and ECM composition. Electroencephalogram recordings revealed Bassoon deletion at excitatory or inhibitory synapses having diverse effects on epilepsy-related phenotypes. While constitutive *Bsn* mutants and GABAergic neuron-specific knockouts (*Bsn^Dlx5/6^cKO*) displayed severe epilepsy with more and stronger seizures than kainate-injected animals, mutants lacking Bassoon solely in excitatory forebrain neurons (*Bsn^Emx1^cKO*) showed only mild impairments. By semiquantitative immunoblotting and immunohistochemistry we show model-specific patterns of neural ECM remodeling, and we also demonstrate significant upregulation of the ECM receptor CD44 in null and *Bsn^Dlx5/6^cKO* mutants. ECM-associated WFA-binding chondroitin sulfates were strongly augmented in seizure models. Strikingly, Brevican, Neurocan, Aggrecan and link protein Hapln1 levels reliably predicted seizure properties across models, suggesting a link between ECM state and epileptic phenotype.

## 1. Introduction

Epilepsy is a pathological brain condition characterized by the propensity to have recurrent seizures. An imbalance of excitation and inhibition has been identified as the underlying cause for epileptic seizure generation (Staley, 2015). Under physiological conditions, the excitation-inhibition ratio is tightly controlled by manifold synaptic and network mechanisms involving also the perisynaptic extracellular matrix (ECM) and its condensed specialization, the perineuronal nets (PNNs) (Chaunsali et al., 2021; Dzyubenko et al., 2021; Mukhopadhyay et al 2021; Saghatelyan et al., 2001). This ECM is rich in hyaluronic acid (HA), chondroitin sulfate proteoglycans, link proteins, and tenascins and controls tissue properties like volume transmission, as well as stiffness and pH of the extracellular milieu. Neural ECM specializations compartmentalize the extracellular space, e.g. around synapses, and are a major regulator of synaptic plasticity and intrinsic excitability (Dityatev et al., 2010). There is increasing evidence that alterations in various components of the ECM, such as reduced hyaluronan levels, may cause seizures (Arranz et al., 2014; Perkins et al., 2017; Vedunova et al., 2013) or contribute to tissue reorganization during epileptogenesis (Dityatev, 2010; Pitkänen et al., 2014).

The aetiology of the epilepsies can be broadly divided into acquired and genetic with genetic or presumed genetic causes underlying ∼20% of the epilepsies (Balestrini et al., 2021; Dityatev 2010). Epilepsies frequently result from disrupted synaptic function, the synaptopathies (Gatto and Broadie, 2010). Accordingly, mutations in genes driving synaptic processes are over-represented in pathways leading to epilepsy (Fukata & Fukata, 2017; Noebels, 2015; do Canto et al., 2021). Among these, molecular processes of presynaptic vesicle cycling and neurotransmitter release frequently result in epilepsy (John et al., 2021; Verhage and Sorensen, 2020).

One of the synaptic proteins implicated in epileptogenic pathways is Bassoon, a presynaptic scaffolding protein that is involved in the assembly and molecular organization of the apparatus for neurotransmitter release in excitatory and inhibitory presynapses (Gundelfinger and Fejtova, 2012; Gundelfinger et al., 2016). Mutations in the murine *Bsn* gene, which result in deletions of large parts of the protein (*Bsn^ΔEx4/5^*), cause severe, rapidly generalizing seizures (Altrock et al., 2003). Epilepsy in these animals is associated with prolonged brain growth and increased levels of brain-derived neurotrophic factor (BDNF) in the forebrain (Angenstein et al., 2007, 2008; Heyden et al., 2011), and abnormal synaptic function and plasticity in the hippocampus and the striatum (Altrock et al., 2003; Ghiglieri et al., 2009, 2010; Sgobio et al., 2010). The epilepsy can be ameliorated by long-term treatment with the anticonvulsant valproic acid (Ghiglieri et al., 2010; Sgobio et al., 2010) and by rearing mutant mice in enriched environment (Morelli et al., 2014). In humans, manifold evidence has been reported that mutations in the *BSN* gene are linked to brain dysfunctions associated with epilepsies. These include intellectual disability (Froukh, 2017), sudden infanticide (Brohus et al., 2021), the Landau-Kleffner syndrome, a rare childhood epileptic encephalopathy (Conroy et al., 2014), and febrile seizures (Skotte et al., 2022). Another very recent study identified multiple variants of the *BSN* gene – both truncations and compound heterozygous mutations – associated with early developmental seizures, thus placing *BSN* among the potential human epilepsy genes (Ye et. al., 2022). However, the underlying molecular mechanism of Bassoon causing these syndromes is still unclear, as is the synapse type-specific contribution of Bassoon deficiency. For studying seizure generation and epileptogenesis, various animal models have been developed (Kandratavicius et al., 2014), among which systemic or intrahippocampal injection of the chemoconvulsant kainate (KA) is frequently used as a model for acquired temporal lobe epilepsy (TLE, Rusina et al., 2021). Unilateral intrahippocampal KA injection produces recurrent seizures, usually evokes unilateral hippocampal sclerosis and reproduces human TLE pathology (Bouilleret et al., 1999; Gröticke et al., 2008; Jeffreys et al. 2016). Thus, in our study we used this model for comparison and as reference for our genetic models.

To further elucidate the significance of Bassoon mutations and the contribution of the neural ECM in epileptic phenotypes, here, we address the following questions: (i) What role does absence of Bassoon from excitatory and inhibitory synapses play in the development of epilepsy in mice? (ii) How does Bassoon-dependent epilepsy compare to the intrahippocampal KA model of TLE? (iii) How do the different types of epilepsy affect the composition of the neural hyaluronan-based ECM and, more generally, the distribution and proteolytic processing of ECM molecules or the hyaluronan receptor CD44 in different epilepsy-relevant brain subregions and subcellular protein fractions/ compartments?

## 2. Material and methods

### 2.1. Mouse models

Mice were housed under standard conditions on a 12-12hrs light-dark cycle. Food and water were provided ad libitum. All *Bassoon* mutant lines were bred in a C57BL/6N genetic background at the LIN. Three lines (*Bsn^ΔEx4/5^*, *Bsn^gt^, Bsn^KO^*) are constitutive and affect the *Bsn* gene in all cells. In Bsn^lx/lx^ the *Bsn* gene is floxed; *Bsn^Emx1^ and Bsn^Dlx5/6^* are conditional knockouts (cKOs) with functional inactivation of the *Bsn* gene in the glutamatergic forebrain and GABAergic interneurons, respectively (for details see Supplementary Fig. S1; Altrock et al, 2003; Annamneedi et al., 2018; Hallermann et al., 2010; Schattling et al., 2019). The animals used for the KA experiments had the C57BL/6J background and were bred under similar specific pathogen-free conditions at the DZNE. Animal breeding and experiments were carried out in accordance with the European Communities Council Directive (2010/63/EU, amendment 2019) and approved by the authorities of the State Saxony-Anhalt (Landesverwaltungsamt Halle) to LIN (TVA No.: 42502-2-1484 LIN) and DZNE (TVA No.: 42502-2-1316 DZNE).

### 2.2 Antibodies, enzymes and drugs

The following antisera/antibodies were used: guinea pig anti-Brevican (diluted 1:1000 for Western blots [WB] and 1:500 for immunohistochemistry [IHC]) and rabbit anti-Brevican (1:1000 for WB) (Seidenbecher et al., 1995; John et al., 2006); rabbit anti-Aggrecan (Millipore, AB1031, 1:500 for WB; RRID:AB_90460); sheep anti-neurocan (R&D Systems, AF5800; 1:200 for WB; RRID:AB_2149717); goat anti-Tenascin-R (Santa-Cruz, sc-9875; 1:200 for WB; RRID:AB_2207007); rabbit anti-Tenascin-C (Cell Signaling Tech, #12221; 1:1000 for WB; RRID:AB_2797849); goat anti-Hapln1 (R&D Systems, AF2608, 1:200 for WB; RRID:AB_2116135); goat anti-Hapln4 (R&D Systems, AF4085; 1:200 for WB; RRID:AB_2116264); rabbit anti-GFAP (Synaptic Systems, 173002; 1:500 for IHC; RRID:AB_887720); sheep anti-CD44 (R&D Systems, AF6127; 1:500 for WB and 1:100 for IHC; RRID:AB_10992919); Wisteria Floribunda Agglutinin (WFA; VectorLab; 1:250 for lectin histochemistry or B1355 biotinylated-WFA antibody, VectorLab 1:500).

Fluorescently labeled secondary antibodies for WB were purchased from Invitrogen (Alexa Fluor 680 & 790, dilution 1:15 000) while antibodies for IHC were obtained from Dianova (Cy3 & Cy5, dilution 1:500). Secondary antibodies coupled to peroxidase (POD) were from Jackson Immuno Research (dilution 1:5 000). Streptavidin was purchased from Invitrogen (Alex Fluor 488 conjugated, S11223, 1:1000). Fluoromount-G (Invitrogen) containing DAPI was used for mounting IHC stained slices. Kainic acid (KA; K0250-10MG) and Chondroitinase ABC (ChABC; C3667) were purchased from Sigma-Aldrich.

### 2.3 Recording of seizure activity

Recordings were done in 3 – 7 months old mice. The timeline of the experimental design for all mouse groups is indicated in Supplementary Fig. S2. EEG recordings were sampled during the 1^st^ week (KA^1wk^) or the 4^th^ week (KA^4wk^) after KA injection.

#### 2.3.1 Surgery and kainate injection

Implantation of electrodes for electroencephalographic (EEG) recordings (electrode E363/96/1.6/Spc, Bilaney consultants GmbH) and cannula placement for kainic acid injection were performed as described previously (Broekaart et al., 2021). After surgery a 3D-printed cap (polylactic acid) composed of two chambers, one for the wireless transmitter and one for the battery, was mounted and the transmitter was attached to the electrode.

For the intrahippocampal kainate model, KA was administered to the CA1 region of the right hippocampus (10 mM, 120 nl at 3 nl/s) (Broekaart et al., 2021). Sham-operated control animals were injected with physiological saline accordingly.

#### 2.3.2 EEG recording and analysis

For recording and analysis experimenters were blinded for genotypes and pharmacological treatment. After a recovery period of about 7-12 days in individual cages, wireless recordings of freely moving animals were performed utilizing the LWDAQ+ software (Open Source Instruments) for up to 10 days, out of which 4-5 consecutive days were sampled (Supplementary Fig. S2). With the LWDAQ Neuroarchiver Tool recordings were cut into 4s frames with a glitch threshold at 200 counts to remove artefacts. In each of the 4s-frames, the ECP19 processor extracted 6 properties from the EEG trace (for details see: Hashemi, 2008-2022, https://www.opensourceinstruments.com/Electronics/A3018/Neuroarchiver.html#Interval%20Processing). The software stored the properties of a selected set of traces collected outside of the actual recording periods for comparison in a library. For *Bsn* mutants, a library of 143 ictal intervals was compiled to be used as references by the software. The library for the KA animals was compiled similarly from their own recordings, with 171 events for one week (KA^1wk^) and 287 events for the four weeks (KA^4wk^) post-KA injection recordings.

The library is then used to classify the remaining recordings. The software computes a space based on the 6 ECP19 processor properties and places each 4s frame in this space before comparing their distance with the closest trace of the library. A 4s frame was considered ictal if the distance to a library point was below an 0.1 threshold distance. It was then confirmed manually whether or not the frames identified as ictal during this process were or not part of an actual seizure. Ictal-like events with a duration below 8s were not considered as seizures. Moreover, the precision of the library was tested against two animals fully reviewed by eye, for each library. We took one animal without seizures to make sure that it was not a false negative and one animal with seizures to make sure that all possible seizures were properly detected by the library. The library was considered accurate once all seizures from the second animal could be selected and it was confirmed that the animal without any detected seizures indeed had none.

Seizure onset patterns can fall into different categories (Perucca et al., 2014). To classify seizure onset types, two observers independently inspected the onset of each seizure and classified them according to pre-ictal and ictal high frequency oscillation analysis. They then compared their classifications and reached consensus. As most seizure onsets fall into the low-voltage fast activity (LVF) group (see results) further events including hypersynchronous and undefined onset types were grouped as ‘others’.

### 2.4 Slice Electrophysiology

#### 2.4.1 Slice preparation

Slice preparation and extracellular field recordings were performed similar to our previous studies (Caliskan et al., 2015; 2016). Mice (10 to 25 weeks old) were decapitated under deep isoflurane anaesthesia and brains were extracted. Horizontal ∼400 µm thick brain slices were cut using an angled platform (12° in the fronto-occipital direction) in ice-cold, carbogenated (5% CO_2_ / 95% O_2_) artificial cerebrospinal fluid (aCSF) containing (in mM) 129 NaCl, 21 NaHCO_3_, 3 KCl, 1.6 CaCl_2_, 1.8 MgCl_2_, 1.25 NaH_2_PO_4_ and 10 glucose (pH 7.4, ∼300 mOsm/kg) with a vibrating microtome (Campden Instruments; Model 752) and quickly transferred to an interface chamber perfused with aCSF at 32 ± 0.5 °C (flow rate: 2.0 ± 0.2 ml/min). The slices were cut until three-to-four slices containing transverse sections of ventral-to-mid hippocampus were obtained. Slices were incubated at least for an hour before recordings were started. Recordings were performed blinded for the experimenter.

#### 2.4.2 Sharp Wave-Ripples and Recurrent Epileptiform Discharges

Analog field potential (FP) signals were pre-amplified and low-pass filtered at 3 kHz using a custom-made amplifier. Then, the analog FP signal was digitized at 5 kHz using an analog-to-digital converter (Cambridge Electronic Design, Cambridge, UK) and stored on a computer hard disc for further signal processing and data analysis as previously described (Caliskan et al., 2016). Sharp wave-ripple (SW-R) recordings were obtained with borosilicate glass electrodes filled with aCSF (resistance in aCSF: 1 MΩ) placed at the pyramidal layer of CA3 and CA1 subregions. Three-to-five min data were recorded, and 2 min artifact-free data were extracted as MATLAB files. Custom written MATLAB-scripts were used to analyze distinct properties of SW-R (MathWorks, Natick, MA). SW detection was performed by using the low-pass filtered at 45 Hz (Butterworth, 8th order) with a detection threshold of 2.5 times the standard deviation (SD) of the lowpass-filtered signal and a minimum interval of 80 ms between two subsequent SW. SW events were stored as 125 ms data stretches centered to the maximum of SW. The mean of the data was used to determine the start and the end point of SW. The area of SW was calculated via integrating the area under curve of low-pass-filtered data between the start and end of the SW event. Ripples were detected using band-pass filter at 120-300 Hz (Butterworth, 8th order) and a detection threshold of 3 SD of the band-pass filtered data. Data stretches of 25 ms (15 ms before and 10 ms after the maximum of SW event) were isolated for further analysis. Triple-point-minimax-determination was used to analyze ripple amplitudes. Ripples with higher than 75% amplitude difference between falling and rising component were discarded. Ripple frequencies were calculated using the time interval between two subsequent ripples.

#### 2.4.3 Statistical analysis of electrophysiological data

SigmaPlot for Windows Version 11.0 (Systat software) was used for statistical analysis of in vitro electrophysiological data. Normality test (Shapiro-Wilk Test) and equal variance test were performed before commencing other statistical tests. For hippocampal network oscillations, statistical differences were determined by Student’s two-tailed t-test or Mann-Whitney U test. To determine whether probability to induce recurrent epileptiform discharges was altered, the Fisheŕs exact test was used. Probability values of *P* < 0.05 were considered as statistically significant. Sample sizes are provided in figure captions (*N*: Number of mice; *n*: Number of slices).

### 2.5 Protein biochemistry and immunoblot analysis

After EEG recording animals were euthanized with carbon dioxide. Brains were removed, frozen in liquid nitrogen and stored at -80°C until further use. Brains were thawed in PBS, homogenized and fractionated essentially as described (Carlin et al., 1980; tom Dieck et al., 1998). The following fractions were collected and probed for the presence of ECM components: homogenate (S1), i.e. raw lysate after removing nuclei and cell debris by 1000xg centrifugation; soluble fraction (S2), i.e. supernatant after 13,000xg centrifugation of S1 containing mainly cytosolic and extracellular soluble components; and synaptosomes, enriched for synaptic and perisynaptic proteins (Carlin et al., 1980, tom Dieck et al., 1998). To remove chondroitin sulfate side chains from core proteins of proteoglycans fractions were digested with ChABC in a 1:400 dilution of a 0.1U/μl solution for 90 minutes at 37°C (Seidenbecher et al., 1995) and subjected to semiquantitative immunoblot analyses.

Protein samples (6-20μg/lane) were separated by SDS-PAGE on 2.5-10% or 5-20% gradient gels and 2,2,2-trichloroethanol in-gel staining was used for loading control. Proteins were transferred onto PVDF membranes (Immobilon-FL, 0.45 μm pore size, Millipore) following standard procedures. Immunoblots were developed with specific primary antibodies followed by incubation with peroxidase-coupled secondary or fluorescent secondary antibodies. Peroxidase-labelled membranes were developed with an ECL Chemocam Imager (INTAS Science Imaging Instruments GmbH) the fluorescently labelled blots were scanned with an Odyssey scanner (LI-COR® Biosciences). The quantification was done with ImageJ (https://imagej.nih.gov/ij/) and values were normalized to samples from respective control animals.

### 2.6 Immunohistochemistry

For immunohistochemistry animals were euthanized with carbon dioxide and after death quickly perfused using a peristaltic pump. Brains were removed and processed essentially as described earlier (Annamneedi et al., 2018). Before any analysis, the negative control sections (without primary antibody incubations) were checked for the absence of fluorescence signal under a fluorescence microscope (Zeiss, Axioplan 2). In case there was a signal, the corresponding channel was disregarded for qualitative and quantitative analysis.

Stained brain slices were observed under a confocal microscope (TCS SP5, Leica Microsystems). Images were taken at 1024 x 1024 px resolution with 15% overlap to ensure proper image stitching over a Z-stack of 17 images 0.59μm apart (9.399μm in total). Images were stitched together using Mosaic-Merge function in LAS X software (Leica Microsystems) and analyzed utilizing the FIJI program package (Schindelin et al., 2012).

### 2.7 Further statistical analyses

#### 2.7.1 Survival analysis

Animals were genotyped after weaning, i.e. about 3 weeks postnatally. A survival analysis was conducted with the data of the five *Bsn* mutant lines and a group of control animals including WT and *Bsn^lx/lx^* animals from the LIN records. It comprised the records of all the animals genotyped between January 1, 2017 to August 1, 2020. The Kaplan-Meier plot and the Cox proportional hazard models are fitted with “spontaneous deaths” only considered as the event. Any other type of death due to experimental use led to a censored data point at the appropriate time. Events before weaning were not counted. In order to estimate the number of animals lost before genotyping, the number of animals of each genotype was determined and compared to the expected Mendelian numbers of the corresponding breedings.

#### 2.7.2 Statistical analysis of ECM expression and correlation

Unless otherwise stated, the statistical analysis was conducted with the R statistical software (https://www.rproject.org/), using the following packages: broom, car, corrplot, dplyr, edfReader, emmeans, ggfortify, glmnet, ggplot2, reshape2, plyr, psych, scales, survminer, tidyr.

Linear multivariate discriminant and regression analyses were performed using XLSTAT (version 2020.5.1, Addinsoft). Corrected R^2^, which corresponds to the fraction of the dependent variable variance that is explained by the linear model, was used as a standard measure to report the regression quality. For regression analysis, minimal set of best predictors was selected, which corresponded to the local maximum of corrected R^2^.

Several statistical tests, reported along with the corresponding p-value, were used for group comparison. Post-hoc tests were carried out only on results of significant ANOVAs, with personalized contrasts on least-square means. Following the post-hoc tests, the *P*-values were corrected for multiple comparisons. The statistical tests were considered significant if *P* < 0.05. For multiple testing, the *P*-value was adjusted with the false discovery rate method (fdr).

## 3. Results

### 3.1. Bassoon mutant mice as an early-onset epilepsy model

The first mouse mutant generated for the *Bsn* gene, *Bsn^ΔEx4/5^*, had a shorter life expectancy due to epileptic seizures in the homozygotes (Altrock et al., 2003). As this mutant still expressed parts of the Bassoon protein, it was not clear, whether the seizures were due to loss of Bassoon function in the CNS or due to gain of function due to the residual 180-kD protein fragment. To assess Bassoon function in the mammalian brain in more depth, additional constitutive and conditional mutants were generated (Supplementary Fig. S1). These include a gene trap line, *Bsn^gt^* (Hallermann et al., 2010) and a conditional line, in which the second protein-coding exon of the *Bsn* gene was flanked by loxP sites (*Bsn^lx/lx^*) and which does not show any overt phenotype (Annamneedi et al., 2018). *Bsn^lx/lx^* mice were used to generate constitutive null mutant mice upon somatic cre-recombination and germline transmission of the *Bsn* gene (*Bsn^KO^*; Schattling et al., 2019) as well as to produce conditional knockouts for Bassoon in glutamatergic forebrain neurons (*Bsn^Emx1^* cKO; Annamneedi et al., 2018) and in GABAergic inhibitory neurons of the murine brain (*Bsn^Dlx5/6^* cKO; Supplementary Fig. S1).

First, we examined survival rates and seizure frequencies of the different mutants. To this end, genotype frequencies of about 3 weeks-old mice, i.e. after weaning, were determined (Fig. 1A). With the exception of *Bsn^Dlx5/6^*, all lines have significantly lower numbers of homozygous mutants than expected as compared to wild-type or *Bsn^lx/lx^* control mice. Currently it is unclear whether this is entirely due to pre- or early postnatal lethality or if, and to which extent, reduced fertility of mutant germ cells contributes to this imbalance of Mendelian ratios. To further assess lethality of the *Bsn* mutation, we monitored survival over the course of 6 months after genotyping by applying Kaplan-Meier estimation (Fig. 1B). From the constitutive mutant lines (*Bsn^ΔEx4/5^, Bsn^gt^, Bsn^KO^*) about 50% of homozygous animals died spontaneously during this period, most of them displaying the typical posture of animals that have died from epileptic seizures. This is consistent with the survival rate initially observed for *Bsn^ΔEx4/5^* (Altrock et al., 2003). *Bsn^Emx1^* cKO mice showed a survival expectancy similar to that of wild types, while *Bsn^Dlx5/6^* cKO animals had a survival rate of about 70%. Long-term EEG recording of a sample of mice revealed that 100% of constitutive mutant mice and of GABAergic cKOs experienced epileptic seizures while only 2 out of 5 of the glutamatergic cKOs displayed seizure activity. Control littermate mice only very rarely suffered from seizures (1 of 26; Fig. 1C). These data suggest that *Bsn^ΔEx4/5^*, *Bsn^gt^*, *Bsn^KO^* and *Bsn^Dlx5/6^* cKO mice may be considered as early-onset epilepsy models, whereas *Bsn^Emx1^* cKO suffer from mild epilepsy.

**Figure 1.**
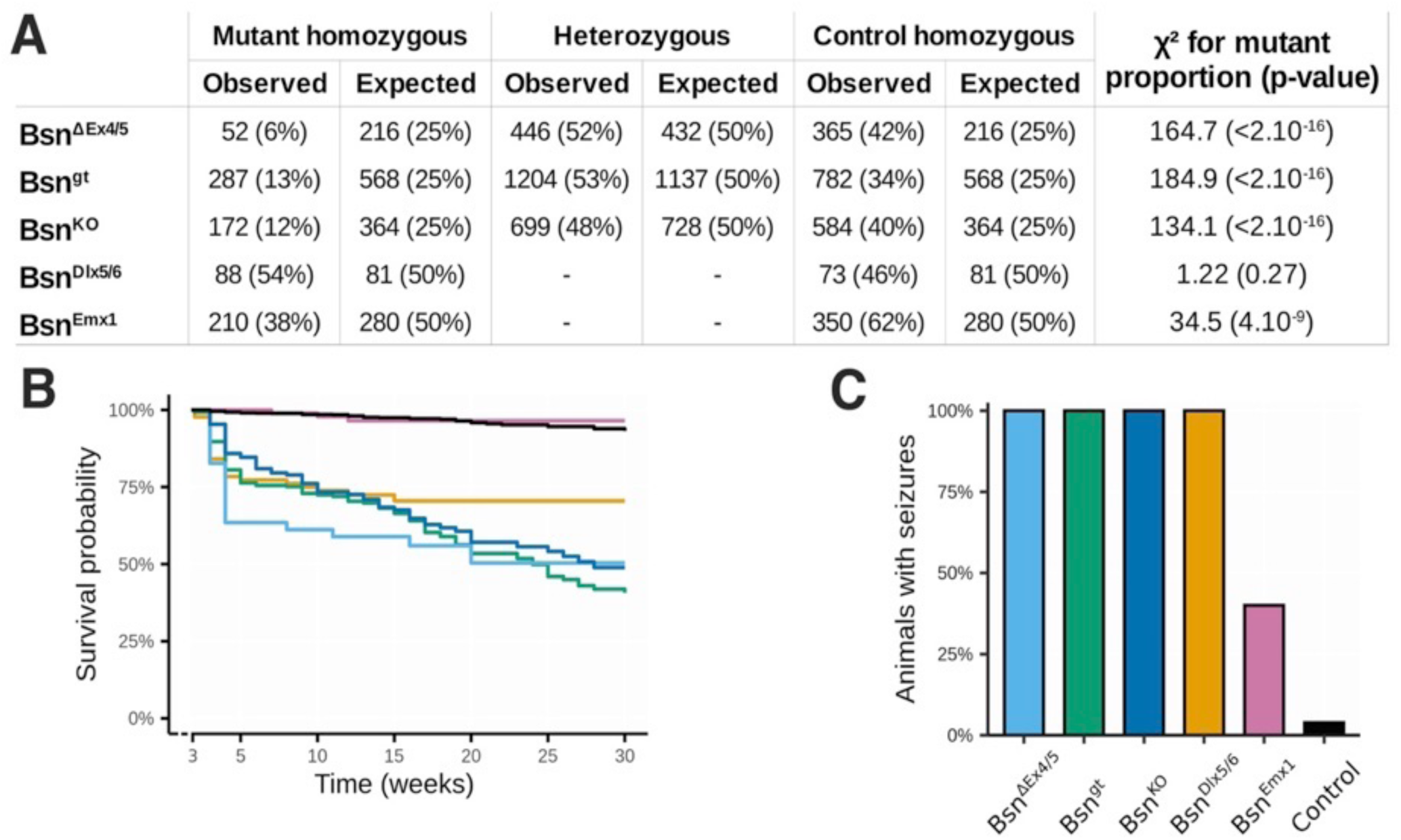
Survival rates and seizure frequencies of Bassoon mutant mice. (A) Genotype frequencies of Bassoon-mutant animals after weaning (3 weeks-old) as compared to expected Mendelian ratios. With the exception of *Bsn^Dlx5/6^*, all the lines have lower numbers of homozygous mutants than expected. Chi-squared test was used to assess statistical significance. (B) Kaplan-Meier survival curve for spontaneous death in genotyped Bassoon mutants. Animal numbers included in the analysis were: *Bsn^ΔEx4/5^ n* = 52; *Bsn^gt^ n* = 287; *Bsn^KO^ n* = 172; *Bsn^Dlx5/6^ n* = 88; *Bsn^Emx1^ n* = 210; WT/*Bsn^lx/lx^* as controls *n* = 2154. No sex differences in the survival were observed in the Cox model (*P* = 0.76). (C) Rate of animals of Bassoon mutant lines developing electrographic seizures during an observation period of 6-10 days (*n* = 4-5 per mutant group, *n* = 26 for control animals). Color code in panel B is the same as in C.

### 3.2. Seizure frequencies and types in Bsn mutant vs. intrahippocampal kainic acid-injected mice

Next, we assessed frequency and severity of seizures in *Bsn* mutants and compared them to an established mouse model of TLE following the unilateral injection of kainic acid into the hippocampus (intrahippocampal KA mouse model; Broekaart et al., 2021). Examples of telemetric EEG recordings of the various mouse models are given in Figure 2A. Seizure frequencies varied from several events per day to one event about every three days. The highest seizure rate with more than four (4.19 ± 0.59) events per day was observed for the *Bsn^KO^* mice. The other constitutive mutants *Bsn^gt^* (2.12 ± 0.45) and *Bsn^ΔEx4/5^* (1.09 ± 0.52) as well as the *Bsn^Dlx5/6^* (1.18 ± 0.39) cKO line lacking Bassoon in inhibitory neurons also experienced on average more than one seizure event daily. In contrast, the *Bsn^Emx1^* cKO line lacking Bassoon expression in excitatory forebrain neurons displays significantly lower seizure rates; here, two out of 5 mice tested have epileptic seizure rates of 0.5-1 daily (Fig. 2B). For comparison, we examined mice of the intrahippocampal KA model one week (KA^1wk^) and four weeks (KA^4wk^) after unilateral kainate injection. Here, in 5 of 12 animals of the KA^1wk^ group and in 5 of 9 mice of the KA^4wk^ group seizure activity was observed during the observation period of 5 consecutive days (Supplementary Fig. S2) with on average 0.54 ± 0.24 seizures per day and group and 0.33 ± 0.14 seizures per day and group, respectively (Fig. 2B). No seizures were detected in control mice for the intrahippocampal KA model, which underwent similar surgery, but received physiological saline injection.

**Figure 2.**
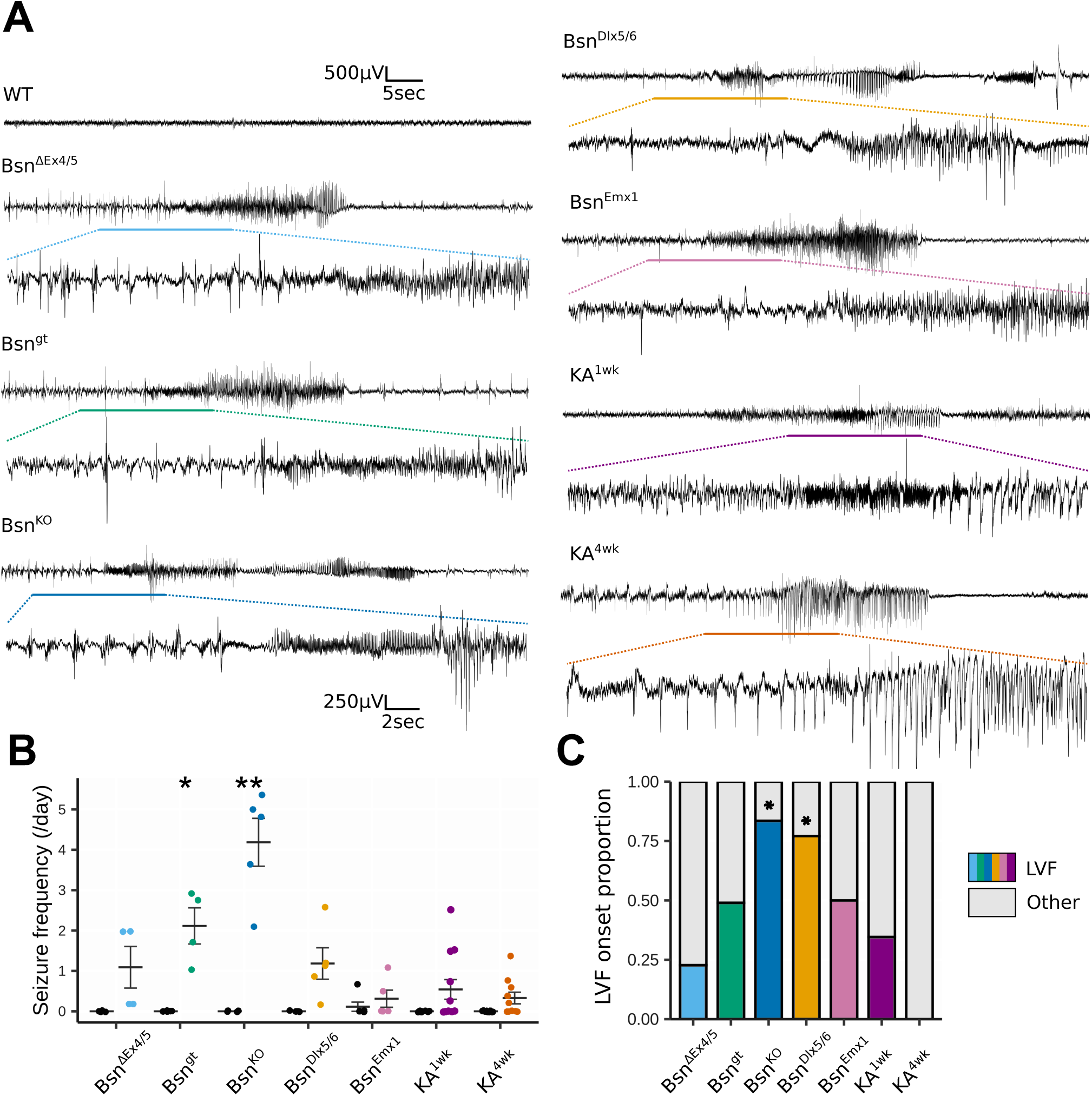
Characterization of seizures in *Bsn* mutant mice as compared to KA-injected animals. (A) Examples of EEG recordings from control animals (*Bsn^lx/lx^*) and each of the *Bsn* mutant lines as well as from intrahippocampally kainate-injected animals, a model of mesial TLE. For each line, a typical example of an epileptic seizure and a zoom on the onset signature is presented. (B) Plot of the average seizure frequency in all *Bsn*-mutant lines and kainate-injected animals both 1 week and 4 weeks post-injection. (C) Proportion of low-voltage fast (LVF) onsets (color shades, color code as in Fig. 1C) over all observed seizures in the different epileptic lines (total events per mouse group: *Bsn^ΔEx4/5^ n* = 22; *Bsn^gt^ n* = 49; *Bsn^KO^ n* = 109; *Bsn^Dlx5/6^ n* = 74; *Bsn^Emx1^ n* = 16; KA^1wk^ *n* = 26; KA^4wk^ *n* = 17). Statistics: * P < 0.05; ** P < 0.01.

Assessment of seizure onset types (Fig. 2A, C) revealed that in particular *Bsn^KO^* and *Bsn^Dlx5/6^* lines display a high proportion of low-voltage fast activity (LVF) onset signatures (84 ± 6% and 75 ± 17%, respectively). Mechanistically, LVF onset seizures are thought to rely on the inhibitory circuits to trigger seizure activity (Elahian et al., 2018). Mixed onset types were observed for other *Bsn* mutants as well as for the intrahippocampal KA model one week after injection (KA^1wk^), whereas in KA^4wk^ animals, other onset types including hypersynchronous onset seizures prevailed (Fig. 2A, C).

### 3.3. Spontaneous recurrent epileptiform discharges in entorhinal-hippocampal slices of Bsn-mutant mice

The Bassoon-mutant line *Bsn^ΔEx4/5^* has been characterized by the occurrence of interictal spikes in the cortex and hippocampus (Altrock et al., 2003). The appearance of hippocampal sharp waves (SW) and the co-occurrence of spontaneous recurrent epileptiform discharges (REDs) can provide hints to epileptic events as already minor changes in hippocampal circuitry (e.g., inhibitory tonus) can transform SW-Rs into pathological epileptiform activity (Behrens et al., 2007; Buzsaki, 2015; Cheah et al., 2021; Karlocai et al., 2014; Liotta et al., 2011). Therefore, to gain deeper insight into potential seizure-generating mechanisms in *Bsn* mutants, we investigated three mutant lines derived from mice with the floxed *Bsn* gene, i.e. *Bsn^KO^*, *Bsn^Emx1^* and *Bsn^Dlx5/6^* for the occurrence of spontaneous REDs as well as for SWs in the hippocampus (Fig. 3). In *Bsn^KO^* mice, about one quarter (26.0%) of entorhinal-hippocampal slices (7 out of 27 slices; 3 out of 5 mice) showed spontaneous REDs, while no such events were observed in control *Bsn^lx/lx^* mice (Fig. 3A, B). The SW incidence was similar in CA3 (*t*_(41)_ = -0.0816, *P* = 0.935), but significantly reduced in CA1 (*t*_(57)_ = 3.404, *P* = 0.001) of *Bsn^KO^* as compared to control mice (Fig. 3A, C). SW area in CA1 (Mann–Whitney *U* = 247, *P* = 0.019) and CA3 (Mann–Whitney *U* = 109, *P* = 0.007) and ripple amplitude in CA3 (Mann–Whitney *U* = 95, *P* = 0.002) were significantly augmented in the null mutants as compared to controls (Supplementary Fig. S3).

**Figure 3.**
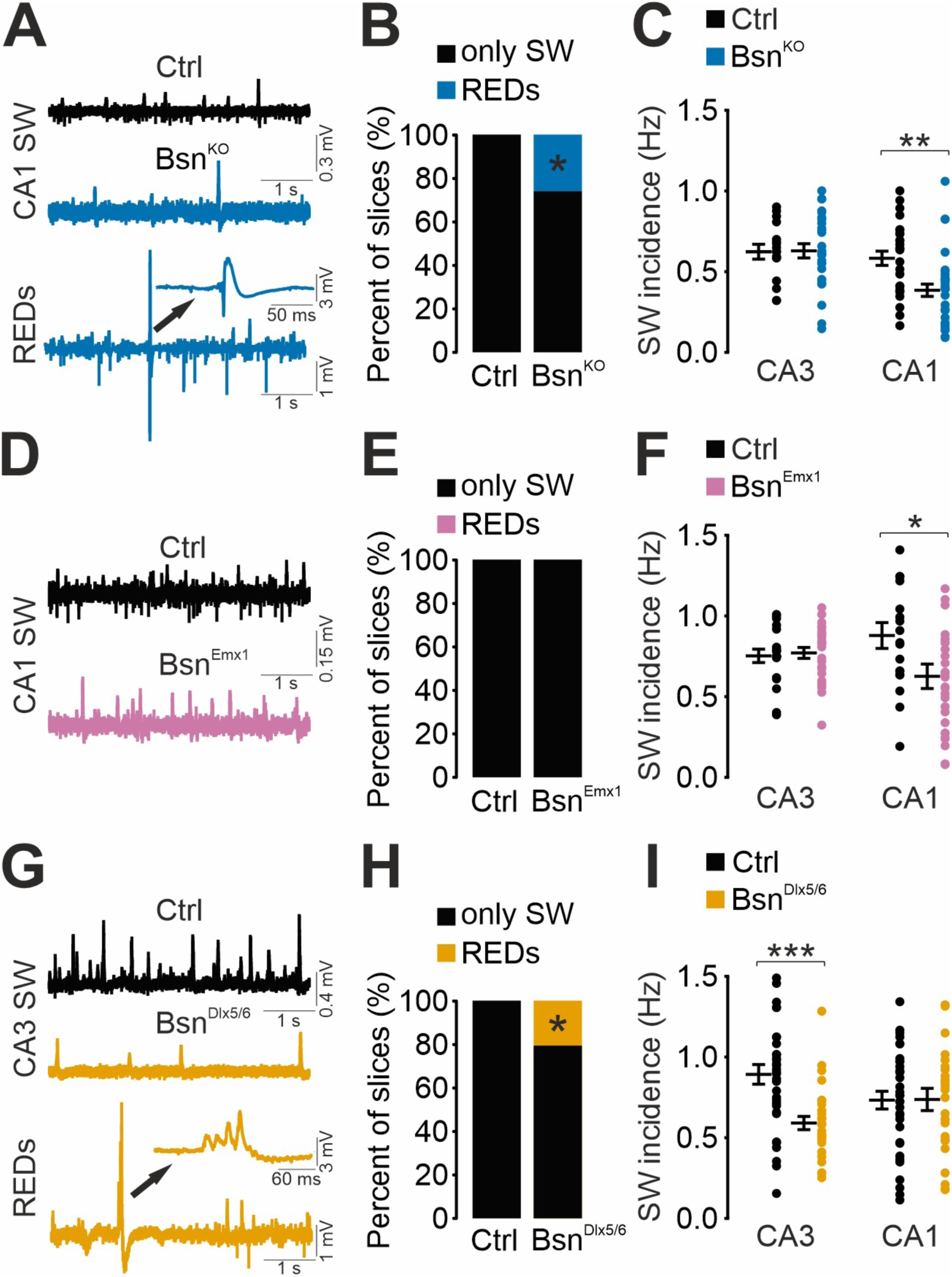
Spontaneous recurrent epileptiform discharges (REDs) and sharp wave (SW) incidences in entorhinal-hippocampal slices of *Bsn*-mutant mice. **A)** Representative field potential traces illustrating reduced sharp wave (SW) incidence in hippocampal CA1 of Bsn^KO^ mice. Note the co-presence of spontaneous REDs in slices from Bsn^KO^ mice. **B)** About 25% of entorhinal-hippocampal slices of Bsn^KO^ mice (7 out of 27 slices; 3 out of 5 mice) exhibited REDs in addition to SW while no REDs were evident in slices from control (Ctrl = *Bsn^lx/lx^*) mice (0 out of 16 slices from 5 mice). **C)** Summary graph illustrating reduced incidence of SW events in CA1 of *Bsn^KO^* mice. **D)** Representative field potential traces illustrating a mild reduction in the SW incidence of *Bsn^Emx1^* mice. **E)** Slices obtained from Ctrl mice (0 out of 19 slices from 5 mice) and *Bsn^Emx1^* mice (0 out of 26 slices from 5 mice) exhibited no spontaneous REDs. **F)** Summary graph illustrating reduced incidence of SW events in the hippocampal CA1 of *Bsn^Emx1^* mice. **G)** Representative field potential traces illustrating reduced SW incidence in the hippocampal CA3 of *Bsn^Dlx5/6^* mice. Note the co-presence of spontaneous REDs in slices from *Bsn^Dlx5/6^* mice. **H)** About 20% of entorhinal-hippocampal slices of *Bsn^Dlx5/6^* mice (7 out of 34; slices; 4 out of 10 mice) exhibited REDs in addition to SW. No REDs were observed in Ctrl mice (0 out of 33 slices from 8 mice). **I)** Summary graph illustrating reduced incidence of SW events in the hippocampal CA3 of *Bsn^Dlx5/6^* mice. Data in **C**, **F** and **I** are presented as mean ± SEM. Data in **B**, **E** and **H** are presented as percent of number of slices included in the study. **B, E, H:** Fisheŕs exact test; **C, F:** Student’s two-tailed test; **I: CA3:** Mann-Whitney U test; **CA1:** Student’s two-tailed test. \**P* < 0.05, \*\**P* < 0.01, \*\*\**P* < 0.001.

In slices from cKO mice lacking Bassoon in excitatory forebrain neurons, no REDs were observed, while the SW incidence was significantly reduced in CA1 (*t*_(45)_ = 2.234, *P* = 0.030), but not in CA3 (*t*_(43)_ = -0.337, *P* = 0.738) (Fig. 3D-F). The SW area (Mann–Whitney *U* = 136, *P* = 0.011) and ripple amplitudes (Mann–Whitney *U* = 153, *P* = 0.032) were augmented and a slight increase in ripple frequency (*t*_(43)_ = 2.377, *P* = 0.022) was detectable in hippocampal CA3 of *Bsn^Emx1^* mice (Supplementary Fig. S4).

In contrast, about one fifth of slices (20.5%; 7 out of 34 slices; 4 out of 10 mice) from cKO mice lacking Bassoon in GABAergic interneurons displayed REDs (Fig. 3G, H). A clearly reduced SW incidence was detectable in the CA3 (Mann–Whitney *U* = 212.5, *P* < 0.001), but not the CA1 region (*t*_(59)_ = -0.0494, *P* = 0.961) of the *Bsn^Dlx5/6^* mice as compared to controls (Fig. 3G, I). In addition, increased ripple frequencies were detected in these mice (CA3: *t*_(60)_ = -2.904, *P* = 0.005; CA1: *t*_(59)_ = -2.086, *P* = 0.041) (Supplementary Fig. S5).

These data disclose an increased propensity for the generation of spontaneous REDs in entorhinal-hippocampal slices of *Bsn^KO^* and *Bsn^Dlx5/6^* mice consistent with the observed severity of epileptic phenotype developed by these mice as compared to the *Bsn^Emx1^* line, and suggest that disturbed interneuron functions strongly contribute to the epileptic phenotype.

### 3.4 Epilepsy-related changes in the ECM in subcellular protein fractions

The ECM surrounding neurons is modified following seizures (Dityatev 2010). Therefore, we next aimed to compare epilepsy-related changes in the hyaluronan-based ECM to search for correlations between epilepsy phenotype characteristics and altered ECM patterns in the different mouse groups. How do seizure severity and frequency affect ECM composition? The HA-based matrix is strongly enriched in PNNs, but is also widely distributed in the brain parenchyma filling the entire extracellular space and in particular wrapping synaptic junctions (Dityatev et al., 2010). To study the alterations in the ECM in the different epilepsy models, we collected the forebrains of animals after the EEG recordings (for the experimental schedule see Supplementary Fig. S2) and isolated subcellular fractions. The homogenate fraction (Hom) was obtained after removal of debris, the soluble fraction (Sol) contains intracellular and extracellular proteins not associated with macromolecular complexes, membranes, and organelles, and the synaptosomal fraction (Syn) comprises synaptic and perisynaptic material. Indeed, we found the ECM composition to be differentially regulated in our epilepsy models with some regulations being specific to the soluble or synaptosomal subcellular fractions, indicating particular changes in solubility or synapse association (Fig. 4). In the brain samples of weakly epileptic Bsn^Emx1^ mice, we did not observe any difference to controls. In the samples of the KA-treated animals, only one statistically significant change was found for Brevican levels, which were reduced in KA^4wk^ homogenates (Fig. 4Aa). In the epileptic Bsn mouse models, a distinct pattern of ECM regulation evolved, with clear upregulation of Aggrecan and Hapln4 in the homogenates of Bsn^Dlx5/6^ and Bsn^gt^ mice, respectively (Fig. 4Aa-c). Together with 130kDa neurocan fragment and with the link protein Hapln1, Aggrecan and its major cleavage product were also upregulated in the soluble fractions (Fig. 4Ba). Finally, in the synaptosomal fractions strongest effects were found for Hapln4, 250kDa Aggrecan and 130kDa Neurocan, all being enriched in Bsn-deficient epileptic mice (Fig. 4Ca). It is noticeable that full-length Brevican clearly stands out because this is the only significantly downregulated molecule among all investigated candidates (Fig. 4Aa, Ba-c, Ca).

**Figure 4.**
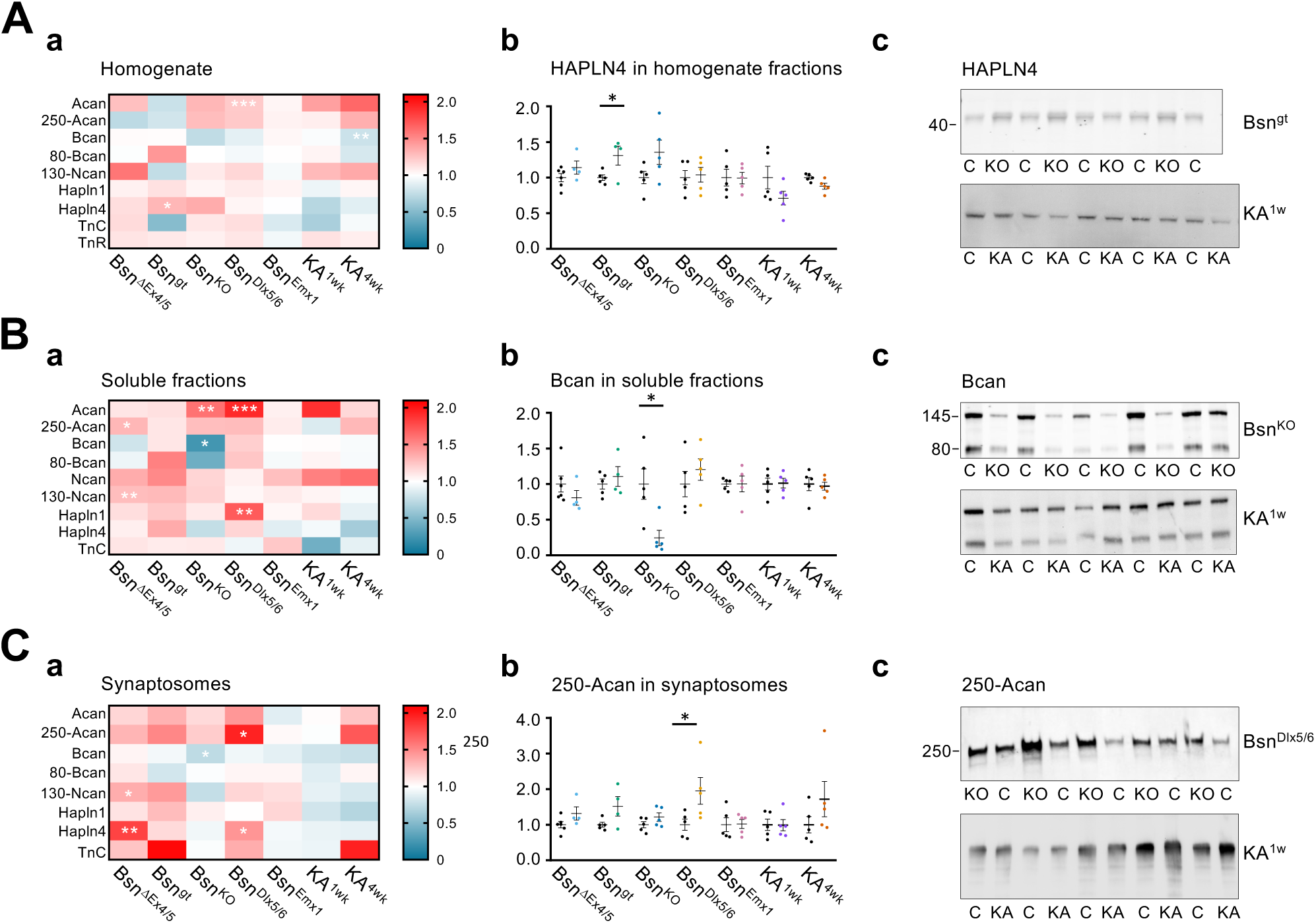
Differences in the signature of components of the hyaluronan-based neural ECM in subcellular forebrain fractions from the different epilepsy models. **(A)** homogenate fractions; **(B)** soluble fractions; **(C)** synaptosomal fractions. Subpanels **(a)** show heatmaps indicating the levels of all ECM components investigated in the homogenates **(A.c)**, soluble fractions **(B.c),** and synaptosomes **(C.c)** of all animal groups, relative to their respective controls. Significant changes are marked with asterisks. Color bar indicates protein level changes. Subpanels **(b)** show comparisons of the levels of Hapln4 in the homogenate fraction **(A.b)**, of full-length Brevican in the soluble fraction **(B.b)** and of 250kDa Aggrecan in the synaptosomal fraction **(C.b)** in mutants to their respective controls. Subpanels **(c) s**how examples of original immunoblots for Hapln4 in the homogenate of Bsn^gt^ and KA^1w^ mice **(A.c)**, for Brevican in the soluble fractions of Bsn^KO^ and KA^1w^ mice **(B.c)**, and for synaptosomal 250kDa Aggrecan fragment in Bsn^Dlx5/6^ and KA^1w^ mice **(C.c)**. Molecular mass of bands is indicated (in kDa). (Statistics: Student’s or Welch’s t-test, or Mann-Whitley rank sum test depending on equal variance and normality of compared samples; *P < 0.05; ** P < 0.01; *** P < 0.001).

In the epileptic Bsn^gt^ line we detected several differences but only the upregulation of Hapln4 survived statistical testing (Fig. 4Aa), while differences in the other investigated molecules did not reach significance, potentially due to higher within-group variability in this strain.

Increased neuronal activity and plasticity reportedly can induce cleavage of ECM components (Dityatev et al., 2007; Mitlöhner et al., 2020; Valenzuela et al., 2014). Interestingly, the ratio between 80kDa cleaved and total Brevican was significantly increased in the homogenate fractions of the highly epileptic *Bsn^gt^* mutants and also in the soluble and synaptosomal fractions of *Bsn^KO^* mice, but not in any of the kainate animal samples. This increase in proteolysis seems to be Brevican-specific, as neither for Aggrecan nor for Neurocan a similar difference in cleavage ratios was observed (Supplementary Fig. S6).

### 3.5 Multivariate analyses of ECM changes in epilepsy models

Correlation analysis revealed a complex pattern of interdependencies between expression and cleavage of ECM proteins in various subcellular fractions and significant correlations between seizure and ECM parameters (Supplementary Fig. S7). To simultaneously consider and represent ECM remodeling in all studied epilepsy models, a linear multivariate discriminant analysis was performed. The differences between epilepsy models were optimally projected in the space of two factors constructed as linear combinations of multiple ECM measures (Fig. 5A). The first factor explaining 47.87% of the total variance was most negatively loaded by expression of cleaved Brevican in soluble and synaptosomal fractions and Tenascin-C in homogenate, and positively loaded by the ratio between the cleaved and total Brevican in soluble and synaptosomal fractions and expression of Hapln4 in the homogenate. The second factor explaining 16.47% of the total variance was most negatively loaded by the ratio of cleaved to total Aggrecan in the homogenate, total Aggrecan in homogenate and soluble Tenascin-C. Expression of cleaved Aggrecan in the homogenate, soluble Aggrecan and cleaved Brevican in synaptosomes provided the highest positive loads to the second factor. Thus, it appears that changes in Brevican and Aggrecan expression in multiple fractions are correspondingly represented by the first versus second factors. The discriminant analysis revealed that (i) most dramatic alterations in ECM measures related to the first factor were observed in *Bsn^KO^* mice; (ii) ECM remodeling in *Bsn^gt^* mice was in the same direction as in *Bsn^KO^* mice but milder; (iii) the second factor revealed differences between *Bsn^Dlx5/6^* and KA^1wk^ mice versus *Bsn^Emx1^*, *Bsn^ΔEx4/5^* and KA^4wk^ mice. This is in line with the analysis of individual ECM measures, which revealed the most distinct ECM remodeling in *Bsn^KO^* and *Bsn^gt^* mice. The difference in the second factor between mice lacking Bassoon in inhibitory versus excitatory neurons may reflect that this factor describes the cleavage and solubilization of Aggrecan, the major component of PNNs associated with fast-spiking interneurons. It is noteworthy that ECM remodeling 1 week after KA injection was more reminiscent to the phenotype of *Bsn^Dlx5/6^* mice with *Bsn* deficiency in inhibitory neurons, while 4 weeks after KA injection mice changed the phenotype to that reminiscent of *Bsn^ΔEx4/5^* mice.

**Figure 5.**
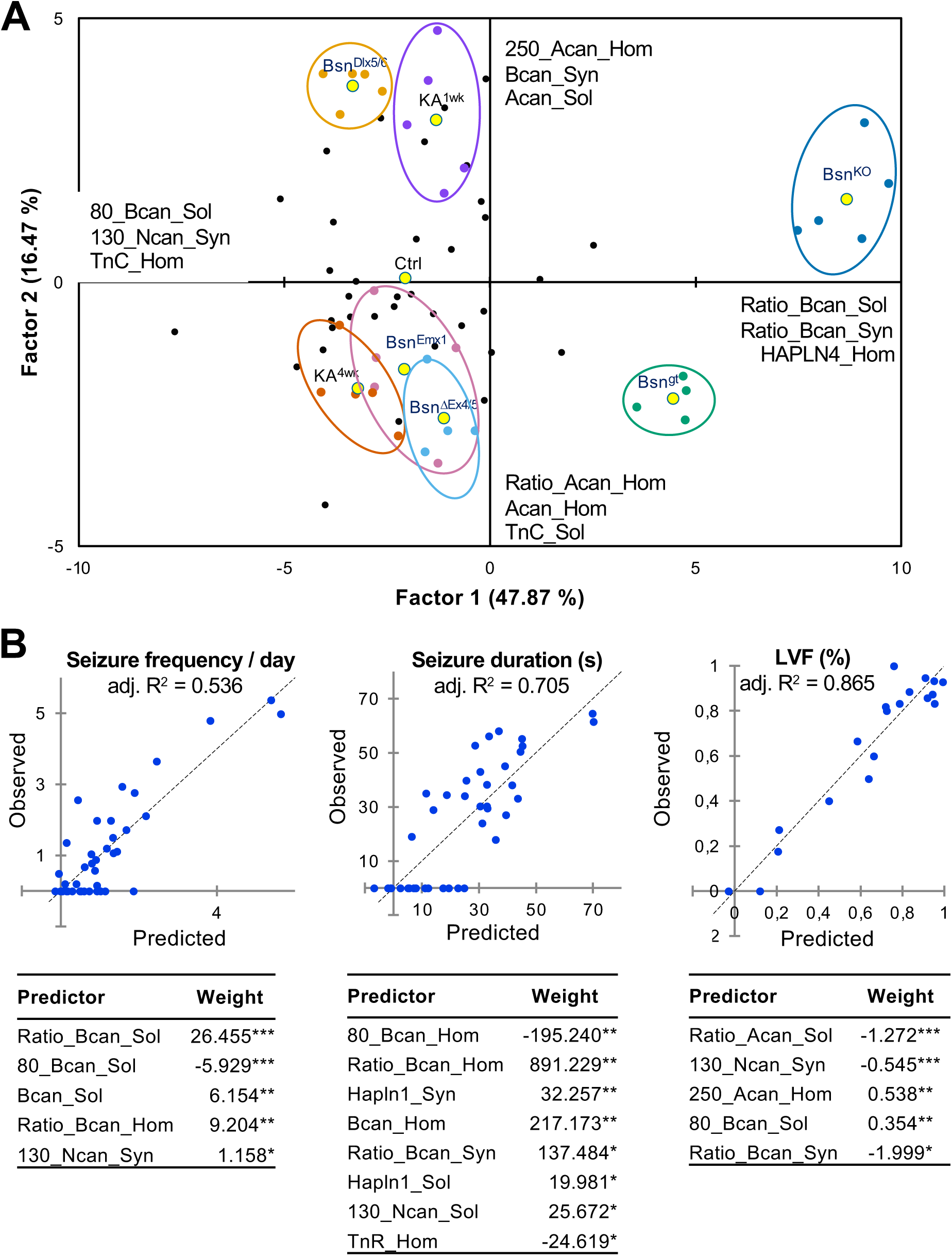
Multivariate analyses of ECM remodeling in mouse models of epilepsy. (A) Linear multivariate discriminant analysis was used to optimally represents differences in the ECM composition between all studied mouse models in the space of two factors constructed as linear combination of studied ECM measures. Three ECM measures most negatively or positively correlating with each of factors are listed next to axes. The ellipsoids outlining data from the same model were added manually to facilitate perception of the relationships between models. (B) Linear multivariate regression analysis was used to determine the ECM measures optimally predicting the frequency and duration of seizures, and proportion of low-voltage fast (%LVF) onsets seizures. Scattergrams depicting the relationships between the observed and predicted seizure parameters are shown. The dashed line is the line of identity. The adjusted R^2^ values provide the fraction of total variance explained by multivariate linear regression model. The best predictors are listed below scatterplots, with their significance: \**P* < 0.05, \*\**P* < 0.01, \*\*\**P* < 0.001.

Multivariate linear regression analysis was used to determine if linear combinations of ECM measures would allow to predict the frequency and duration of seizures, and the fraction of LVF seizures (Fig. 5B). Among significant predictors of seizure frequency and duration were 5 and 8 ECM measures (Fig. 5B) partially overlapping with the first factor revealed by the discriminant analysis and predominantly reflecting Brevican proteolytic cleavage in all studied fractions. The predictors of the fraction of LVF seizures (involving failures in GABAergic transmission) were related to Aggrecan proteolysis and partially overlapped with the second factor. The ECM parameters explained 53.6%, 70.5% and 86.5% of the total variance in the seizure frequency, duration of seizures, and the fraction of LVF seizures, respectively. These data highlight a relevance of ECM remodeling to ictogenesis and/or epileptogenesis.

### 3.6 Spatial distribution of epilepsy-associated ECM changes

To evaluate where in the brain epilepsy-related ECM remodeling is most pronounced, we performed immunohistochemistry in *Bsn^KO^* mice and compared the patterns of WFA labeling and of Brevican immunolocalization with wildtype control patterns (Supplementary Fig. S8). The plant lectin WFA is a widely used indicator for a PNN-typical glycosaminoglycan subtype, which is preferentially linked to Aggrecan (Härtig et al., 2022). We found a wide-spread and evident upregulation of WFA-binding chondroitin sulfates paralleled by an overall loss of Brevican, with both effects being most pronounced in the hippocampus. Therefore, we focused on this brain area for a detailed comparison of our different Bassoon-deficient lines (Fig. 6). We observed Brevican down- and WFA up-regulation strongest in the full knockout and also detectable in the GABAergic neuron-specific knockout *Bsn^Dlx5/6^*, the lines displaying the strongest epileptic phenotypes. Increased WFA signal was concentrated in the granular and molecular layers of the dentate gyrus, but also associated with individual neurons in the CA1 region. Similar upregulation of WFA in the dentate gyrus was found in KA^4wk^ mice (Supplementary Fig. S9).

**Figure 6.**
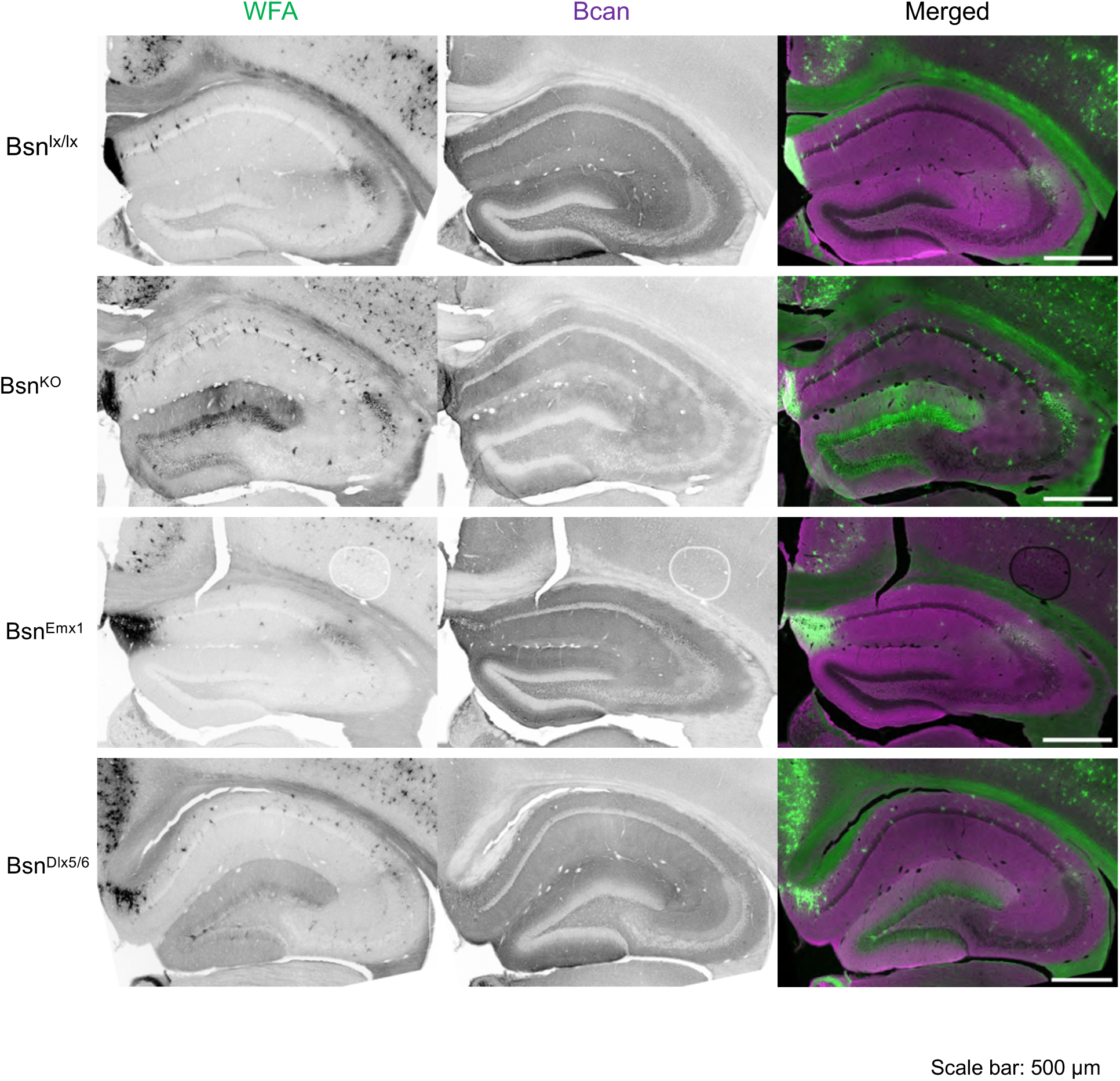
Lectin- and immuno-histochemical distribution of Wisteria floribunda agglutinin (WFA; green) and Brevican (Bcan; purple) in the hippocampus of Bassoon-mutant lines *Bsn^KO^*, *Bsn^Emx1^*, *Bsn^Dlx5/6^*. The floxed mouse *Bsn^lx/lx^* served as control. Scale bar: 500 μm.

### 3.7 The ECM receptor CD44 is upregulated in epileptic mice

In search for a cellular mechanism of disturbed cell-ECM communication we performed a proteomic screen with membrane fractions from brains from *Bsn*-deficient lines *Bsn^KO^*, *Bsn^Emx1^* and *Bsn^Dlx5/6^* and identified the hyaluronan receptor CD44 as a candidate molecule with increased expression in Bassoon full (*Bsn^KO^*) and inhibitory neuron-specific (*Bsn^Dlx5/6^*) knockout mice. With immunoblotting we confirmed a two- to three-fold upregulation of CD44 in the complete as well as in the inhibitory neuron-specific, but not in the weakly epileptic excitatory neuron-specific knockout (Fig. 7). Interestingly, this increase in CD44 immunoreactivity occurred also in the kainate model one week after injection, and it was also detected in the synaptosomal fractions (Fig. 7A-B), suggesting increased perisynaptic ECM binding in epileptic brains. Immunohistochemistry revealed major increase in CD44 immunolabel in the molecular layer of the dentate gyrus as well as in the CA1 *stratum lacunosum moleculare* (Fig. 7C). Double labeling with anti-GFAP antibodies reveals no co-localization, indicating that the increased CD44 levels are unlikely to be produced by astroglial cells (Fig. 7C).

**Figure 7.**
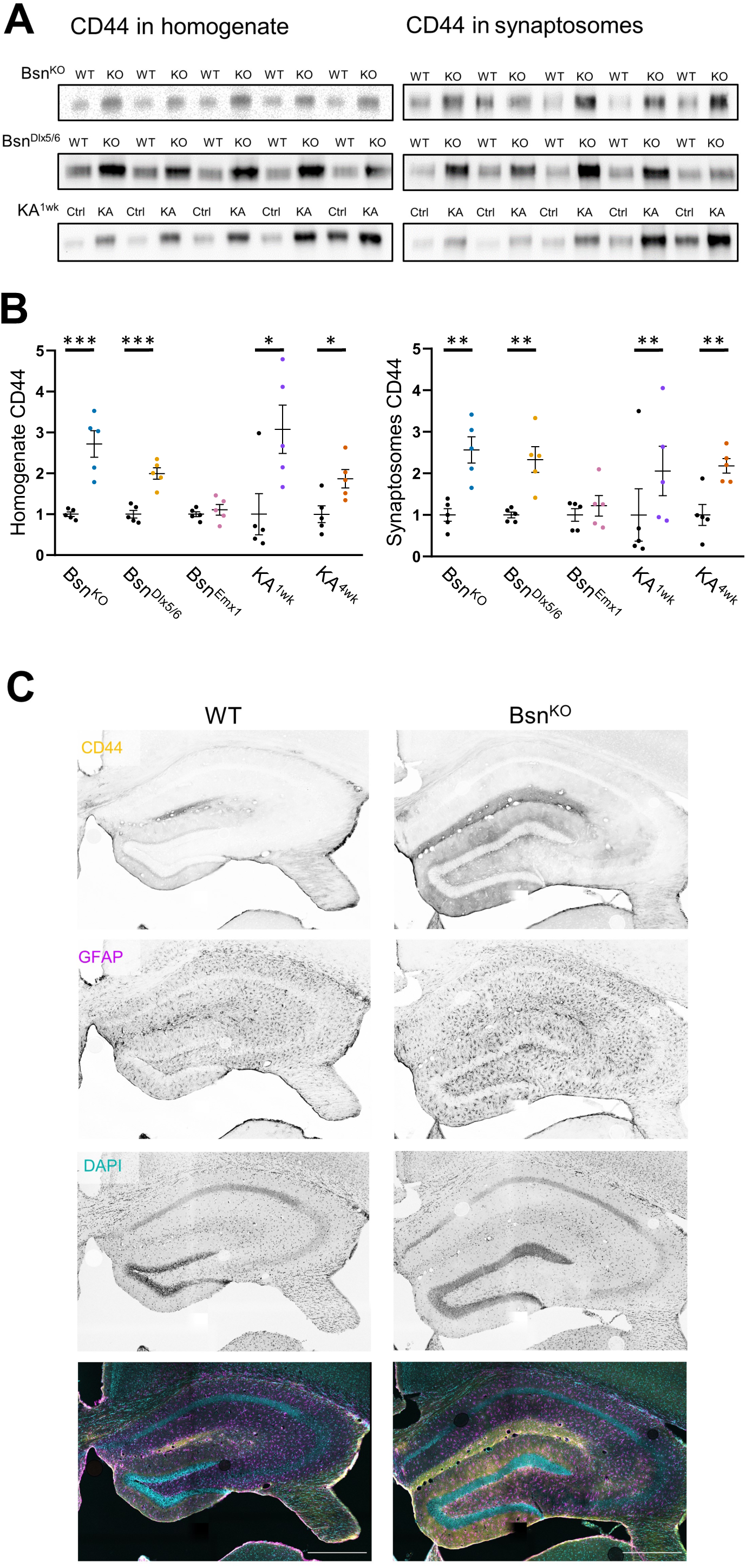
Upregulation of CD44 in epileptic Bassoon mouse lines as well as in the KA model. (A) Immunoblot examples showing CD44 immunoreactivity in the homogenate and synaptosomal fractions of *Bsn^KO^* and *Bsn^Dlx5/6^* as well as KA^1wk^ mouse forebrains. (B) Quantification of CD44 immunoreactivity in the constitutive (*Bsn^KO^*; blue), the excitatory neuron-specific (*Bsn^Emx1^*; purple) and the inhibitory neuron-specific (*Bsn^Dlx5/6^*; ochre) Bassoon-deficient mouse lines as well as in kainate-treated mice (KA^1wk^, violet; KA^4wk^, brown) as compared to their corresponding controls (black dots). (C) Immunohistochemical localization of CD44 immunoreactivity (yellow) in control (WT) and Bassoon-mutant (*Bsn^KO^*) mouse hippocampus. Astrocytes are labeled with anti-GFAP (purple); DAPI staining is shown in cyan. Scale bar: 500μm. Two-way ANOVA with Sidak’s multiple comparisons of KO/KA and WT/Ctrl, \**P* < 0.05, \*\**P* < 0.01, \*\*\**P* < 0.001.

## 4. Discussion

### Bassoon mutants as suitable models for human epilepsy

Initial studies have shown that *Bsn*-mutant mice with partial deletion of the Bassoon coding region affecting all synapses (*Bsn^ΔEx4/5^*) suffer from massive sensory impairment (Dick et al., 2003; Khimich et al., 2005; tom Dieck et al., 2005) and from severe, sometimes fatal epileptic seizures (Altrock et al., 2003). Meanwhile clinical evidence has accumulated that genetic variability in the human *BSN* gene contributes to epilepsies in patients (Conroy et al., 2014; Skotte et al., 2022; Wang et al., 2017; Ye et al., 2022) as well as to other brain disorders, including intellectual disability (Froukh, 2017), bipolar disorder, schizophrenia (Chen et al., 2021), Parkinson’s Disease (Yemni et al., 2021), progressive supranuclear palsy-like syndrome and tauopathies (Martinez et al. 2022; Yabe et al., 2018). Moreover, Bassoon can form intracellular aggregates in multiple sclerosis (Schattling et al., 2019) and aggravate tau seeds in tauopathies (Martinez et al., 2022), suggesting that balanced Bassoon levels are crucial for a healthy brain.

All three constitutive *Bsn* mouse mutants develop severe epilepsy in homozygosity with high seizure-induced lethality (∼50%) between 3 and 30 weeks of postnatal life. Reduced ratios of homozygous animals point to high lethality before genotyping, although reduced fertilization rates by mutated germ cells cannot be excluded. Indeed, viable homozygous mutant animals do not breed. Also, all *Bsn^Dlx5/6^* inhibitory neuron-specific cKO mice develop seizures and display increased lethality between postnatal weeks 3 and 5, while *Bsn^Emx1^* excitatory neuron-specific cKO mice display similar survival rates to wild type control mice and clearly lower seizure incidence.

The majority of patients carrying recently identified *BSN* mutations affecting primarily the C-terminal region of Bassoon (Ye et al., 2022) displayed infancy or childhood onset epilepsies frequently with a history of febrile convulsions. Most of these human mutations seem to have a benign outcome letting the authors speculate that, like in mice, more severe mutations in humans may be fatal (Ye et al., 2022).

A very intriguing finding of this study is that a major contribution to the seizure-inducing pathology originates from Bassoon deletion in GABAergic interneurons. In vivo seizures are paralleled by the occurrence of spontaneous recurrent epileptiform discharges in hippocampal slices of *Bsn^Dlx5/6^* cKO mice, which occur at a similar rate in constitutive knockouts but are absent from mice lacking Bassoon at excitatory forebrain synapses only (Fig. 3). Sharp-wave ripples, complex oscillatory patterns originating from the hippocampus, are considered as hallmarks of episodic memory encoding and behavioral planning (Buzsaki, 2015). As observed either in epileptic *Bsn^Dlx5/6^* or *Bsn^KO^* mice, reduced SW generation and/or emergence of fast ripple oscillations have been consistently demonstrated in other genetic and pharmacologically-induced (e.g., kainic acid) epilepsy models both *in vivo* and *in vitro* (Bragin et al., 2004; Cheah et al., 2019; Engel et al., 2009; Foffani et al., 2007; Jeffreys et al., 2012; Lippman et al., 2022). Furthermore, reduced GABAergic inhibition can lead to pathological conversion of SW-Rs into REDs (Liotta et al., 2011). Indeed, ablation of Bassoon in inhibitory interneurons causes reduced transmission at inhibitory synapses (Annamneedi et al., in prep.), likely causing disinhibition. Earlier studies on *Bsn^ΔEx4/5^* mice that lack functional Bassoon at all chemical brain synapses revealed an abnormal NMDA receptor-dependent short-term potentiation in striatal fast-spiking GABAergic interneurons that was absent in wild-type animals (Ghiglieri et al., 2009) and seems to be triggered via the BDNF/TrkB system (Ghiglieri et al., 2010). Administration of the TrkB antagonist k252a prevented the emergence of this pathological plasticity in fast-spiking interneurons.

What might be the role of the BDNF/TrkB system in epileptogenesis in *Bsn* mutant mice? BDNF levels are massively increased in the cerebral cortex, hippocampus and striatum of *Bsn* mutants (Dieni et al., 2012; Heyden et al., 2011). This increase is paralleled by a steady growth of these brain regions (Angenstein et al., 2007; Heyden et al., 2011). Reportedly, BDNF levels are increased in epilepsy (Binder et al., 2000) and BDNF has crucial effects on interneuron development (Bolton et al, 2000; Ghiglieri et al., 2011; Huang et al., 1999; Marty et al., 1997; Willis et al., 2022). However, whether the increased activity of the BDNF/TrkB system in *Bsn* mice is caused by early developmental epilepsy or due to the absence of Bassoon or actually both is unclear. Our findings from the *Bsn^Emx1^* excitatory neuron cKO mice, which display increased TrkB activity that affects hippocampal development (Annamneedi et al., 2021) but relatively low seizure activity, argue for an at least partially epilepsy-independent effect.

It is well documented that BDNF binds to and forms complexes with glycosaminoglycan components in the neural ECM, like chondroitin sulfates (Kanato et al., 2009), and that these low-affinity binding capacities can be affected in neuropathology (Huynh et al., 2019).

### Epilepsy-associated ECM remodeling

Differences in the expression of ECM components at various stages of epilepsy have been reported (Soleman et al., 2013). For instance, upregulation of chondroitin sulfates and hyaluronic acid was shown in post-mortem hippocampi of mesial TLE patients (Perosa et al., 2002). However, most studies focused on PNNs as most striking neural ECM formations (Chaunsali et al., 2021). Our data show that under epileptic conditions not only the PNNs as largely insoluble condensed ECM structures but also the composition of soluble and perisynaptic ECM are widely affected.

Coming to individual ECM molecules, we discovered correlations between epileptic seizures and the expression Hapln4/Bral2, a link protein which colocalizes with Brevican in PNN (Bekku et al., 2003) and is involved in the regulation of GABAergic synapses (Edamatsu et al., 2018). Surprisingly, Hapln4 regulation is oppositely affected in our genetic and chemical epilepsy models with higher amounts in homogenates and synaptosomal fractions of *Bsn*-deficient mice while KA-injected mice have a tendency for reduced Hapln4 levels, arguing for a differential regulation of this key ECM and PNN component.

The proteoglycan neurocan was already discussed in the literature in other epilepsy models: Okamoto and colleagues (2003) demonstrated a temporary rise of neurocan after systemic KA application causing severe convulsions in rats. Another study also showed a transient increase in mice after intrahippocampal injection of domoate, a kainate-related neurotoxin (Heck et al., 2004). This is in part confirmed by our finding of a significant rise in the 130kDa neurocan fragment, an astrocyte-secreted inhibitor of N-cadherin- and β1-integrin-mediated adhesion and neurite outgrowth (Li et al., 2000).

For Tenascin-R, an increase of immunoreactivity was detected in the hippocampal CA3 neuropil after pilocarpine-induced status epilepticus in mice (Brenneke et al., 2004), whereas here we did not find significant differences between experimental groups. Another study revealed in the pilocarpine model reduced expression of Aggrecan and Hapln1 (McRae et al., 2012), two ECM components, which were upregulated in some of the Bassoon mutants – pointing to obvious mechanistic differences between the pilocarpine and our models.

In the present study, strongest downregulation was observed for the proteoglycan Brevican, one of the most prominent components of the adult perisynaptic and diffuse ECM that is also present in PNNs. This finding is in line with reduced Brevican levels in surgical resections of patients with temporal lobe seizures (Favuzzi et al., 2017). A more recent study reported upregulated Brevican levels in human post-mortem frontal cortex samples from epilepsy patients as compared to controls (Pires et al., 2021), but the post mortem intervals in the control group were more than twice as long as in the epilepsy group, which may cause artificial differences. Finally, our finding of increased Brevican cleavage in Bassoon-deficient epileptic mice corresponds to an earlier report about augmented Brevican proteolysis by ADAMTS in KA-treated rats (Yuan et al., 2002).

In sum, soluble as well as condensed forms of neural ECM become depleted for Brevican but enriched for Aggrecan, WFA-binding glycosaminoglycans and the Neurocan N-terminal fragment. The underlying mechanisms of all these selective ECM reorganization patterns can be manifold and may involve, e.g., seizure-dependent changes in ECM synthesis and posttranslational exocytosis and incorporation into PNN, proteolytic processing, ECM endocytosis and clearance, or even seizure-enhanced autophagy (Otabe et al., 2014), a cellular process which is at presynapses at least in part under the control of Bassoon (Hoffmann-Conaway et al., 2020; Okerlund et al., 2017).

A potential mechanistic hub could be the strongly over-expressed HA receptor CD44, which was already shown to be upregulated under different epileptic conditions, e.g. in KA-treated hippocampal slice cultures (Bausch 2006) and in pilocarpine-injected mice (Borges et al., 2004). CD44 is expressed in synapses and considered to be a modulator of synaptic plasticity (Roszkowska et al., 2016) and reorganizer of neuronal and synaptic circuitry after axon terminal degeneration (Borges et al., 2004).

### Functional implications of seizure-associated ECM remodeling

The strong correlation between seizure parameters and expression of ECM molecules found in this work can be either due to seizure-dependent proteolysis and remodeling of ECM or to modulation of neuronal physiology and synaptic connectivity downstream of ECM remodeling. Evidence that MMP9 knockout or pharmacological inhibition of MMP2/9 strongly suppressed ictal activity and epileptogenesis in different models of epilepsy (Broekaart et al., 2021; Wilczynski et al., 2008) and experiments demonstrating degradation of PNNs by MMP9 under various conditions (Dwir et al., 2020; Stamenkovic et al., 2017; Wen et al., 2018) suggest that MMP9-dependent remodeling of PNNs might determine key aspects of epileptogenesis. Pooling data from several mouse models with genetic and acquired forms of epilepsy, we report here that parameters describing Brevican cleavage in homogenate and soluble fractions are the best predictors of the frequency and duration of seizures. This is in line with studies showing that digestion of PNNs with ChABC or conditional depletion of Brevican expression in PV+ cells impairs excitatory input to these GABAergic interneurons (Favuzzi et al., 2017; Hayani et al., 2018), which could result in insufficient recruitment of PV+ interneurons during bursts of network activity and, eventually, in ictal events. Interestingly, expression of neuronal activity-regulated pentraxin (Narp) is strongly upregulated by seizures and might promote AMPA receptor accumulation in excitatory synapses on PV+ cells (Chang et al., 2010). However, Narp stabilization requires the integrity of perisynaptic ECM of PNNs. When the latter is degraded due to elevated endogenous proteinase activity or enzymatic treatment with hyaluronidase (Vedunova et al., 2013), this powerful endogenous anti-epileptic mechanism can be compromised, resulting in the generation of epileptiform activity. Moreover, a conditional knockout of Aggrecan in PV+ cells ablated PNNs and caused a shift in the population of parvalbumin-expressing inhibitory interneurons toward a high plasticity state (Rowlands et al., 2018). Enzymatic attenuation of Aggrecan-enriched PNNs or depletion of Brevican also lead to increased excitability of PV+ interneurons (Dityatev et al., 2007; Favuzzi et al., 2017; Hayani et al., 2018). These data suggest a complex dependence of synaptic innervation, excitability and plasticity of PV+ cells on the ECM of PNNs. Remodeling of PNNs appeared to be of functional relevance to LVF onset seizures because i) a highly significant prediction of the proportion of such seizures by a linear combination of parameters describing proteolysis of Brevican and Aggrecan, as found in the present study, and ii) previous recordings in rodent models and in human TLE patients, which revealed that seizure-like events with LVF onset are initiated by synchronous inhibitory discharges (Avoli et al., 2016; Elahian et al., 2018).

In addition to Brevican processing, the level of cleaved Neurocan expression in the soluble fraction and expression of Tenascin-R in homogenates (with positive and negative contributions, respectively) proved to be predictors of seizure duration. Corroborating these findings, other studies revealed an increased excitatory input to excitatory neurons and decreased perisomatic inhibition in Tenascin-R-deficient mice (Saghatelyan et al., 2001). These data and the fact that several seizure predictors are related to expression of Brevican, Neurocan, and Hapln1 in synaptosomes suggest that also remodeling of perisynaptic ECM at excitatory synapses on principal cells may be relevant to synaptic remodeling underlying epilepto- and/or ictogenesis. A previous modeling study supports the view that activity-dependent proteolysis of ECM may lead to a switch in the steady state of ECM expression and neuronal activity (Lazarevich et al., 2020). Moreover, proteolysis of ECM molecules has been associated with the key hallmarks of TLE such as granule cell dispersion, mossy fiber sprouting and astroglyosis (Dityatev and Fellin, 2009; Dityatev, 2010). Further studies with brain region- and cell type-specific expression of proteolysis-resistant forms of ECM molecules are warranted to dissect multiple mechanisms by which ECM remodeling leads to formation of hyperexcitable networks.

## Supporting information

Supplemental Figures S1 - S9 with legends

## List of abbreviations

Acan: Aggrecan
Bcan: Brevican
Bsn/BSN: Bassoon gene
BDNF: Brain-Derived Neurotrophic Factor
ChABC: Chondroitinase ABC
ECM: Extracellular Matrix
HA: Hyaluronic Acid
Hapln1: Hyaluronan And Proteoglycan Link Protein 1
Hapln4: Hyaluronan And Proteoglycan Link Protein 4
Hom: Homogenate
KA: Kainic Acid
KO: Knockout
cKO: Conditional knockout
LVF: Low voltage fast activity
Ncan: Neurocan
Pv: Parvalbumin
PNN: Perineuronal Net
RED: recurrent epileptiform discharges
SW/SW-R: Sharp waves/Sharp wave ripples
Sol: Soluble protein fraction
Syn: Synaptosomal protein fraction
TLE: Temporal Lobe Epilepsy
WFA: *Wisteria Floribunda* Agglutinin
WT: Wild type

## Acknowledgements

The authors gratefully acknowledge expert technical support by Kathrin Hartung, Michelle Wirsum, Kathrin Pohlmann, Isabel Herbert and animal caretakers at LIN. We would like to thank Karl-Heinz Smalla for valuable advice on the protein biochemistry. A.B. and S.J. were fellows of the Marie Curie ITN ECMED (Grant agreement ID: 642881). A.B. and A.A. were supported by a LINseeds grant. J.S. is a member of the SynAGE graduate program. Research in the authors’ labs is supported by the Deutsche Forschungsgemeinschaft (362321501/RTG 2413 SynAGE to A.D., E.D.G. and C.I.S. and 425899996/CRC1436 to A.D. and C.I.S.). The work was also supported by grants from Center for Behavioral Brain Sciences (CBBS, promoted by Europäischer Fonds für regionale Entwicklung - EFRE (ZS/2016/04/78113) and CBBS ScienceCampus funded by the Leibniz Association (SAS-2015-LIN-LWC) to G.C., A.A. and C.M.- V.

## Authors’ contributions

E.D.G., A.D. and C.I.S designed the study and wrote the manuscript. A.B. and S.J. performed most of the experiments including immunoblot analysis and EEGs; A.B., S.J., A.D. performed the statistical analyses of the data; G.C. and O.S. performed and analyzed the *in vitro* electrophysiological experiments; A.B., S.J., A.A. and J.S. contributed immunohistochemical data; A.F. designed and established mouse mutants; C.M.-V. organized and supervised mouse genetics and breeding. R.C.W. and M.C.W. advised and contributed to the EEG analyses; all authors contributed to the final version of the manuscript.

