## Supplemental Figures S1 - S9 with legends for "Neural Extracellular Matrix Remodeling Signatures in Genetic and Acquired Mouse Models of Epilepsy"

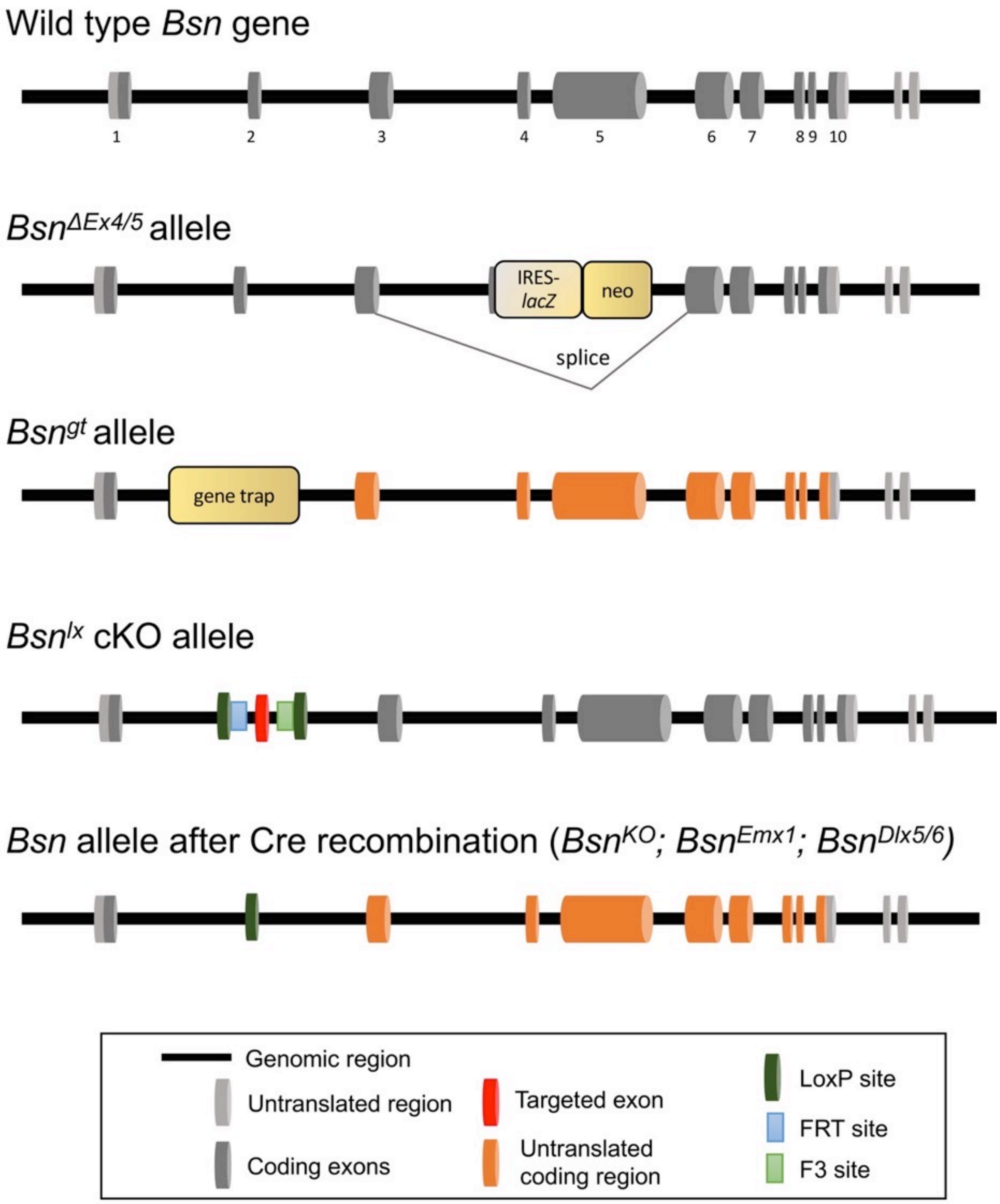


**Figure S1.** Mutant alleles of the *Bsn* gene used in the study. The murine *Bsn* gene includes 10 protein-coding exons. Mice with a constitutive deletion of exons 4 and 5 ($Bsn^{\Delta Ex4/5}$) were first reported by Altrock et al. (2003). Mice with a gene-trapped allele of the *Bsn* gene ($Bsn^{gt}$) were generated from Omnibank ES cell line OST486029 by Lexicon Pharmaceuticals (Hallermann et al., 2010). Mice with floxed *Bsn* gene ($Bsn^{lx/lx}$) were generated for us by Taconic Artemis GmbH (Annamneedi et al., 2018; Schattling et al., 2019)). They behave like wild types. From these mice null mutants ($Bsn^{KO}$) were generated by crossing them with germline Cre-expressing mice (*B6.C-Tg(CMV-cre)1Cgn/J*) (https://www.jax.org/strain/006054). Conditional knockout (cKO) mice were generated by breeding $Bsn^{lx/lx}$ animals with either $Emx1^{tm1(cre)Krj}$ ($Bsn^{Emx1}$) or *Tg(dlx5a-cre)1Mekk* ($Bsn^{Dlx5/6}$) mice. $Bsn^{Emx1}$ cKO mice lack Bassoon expression in glutamatergic neurons of the forebrain (Gorski et al., 2002) and $Bsn^{Dlx5/6}$ cKOs are Bassoon-deficient for GABAergic interneurons (Monory et al., 2006) (http://www.informatics.jax.org/reference/J:122066).

Supplementary Figure S2

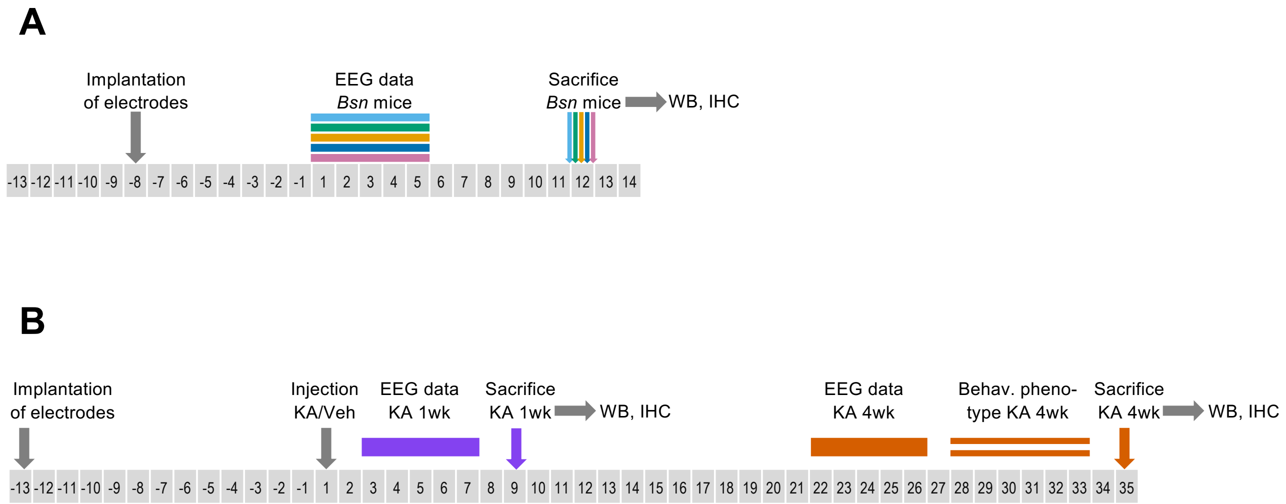


**Figure S2.** Schematic illustration of the experimental design for *Bsn*-mutants (**A**) and kainate-treated animals (**B**). The period of EEG data sampling is indicated. Color code for mutants and treatments as in other figures throughout the manuscript. $KA^{1wk}$ and $KA^{4wk}$, mice from which EEG was sampled during the first week and fourth week, respectively, after kainate injection into the hippocampus. After EEG sampling mice were sacrificed for western blot analysis (WB) and immunohistochemistry (IHC).

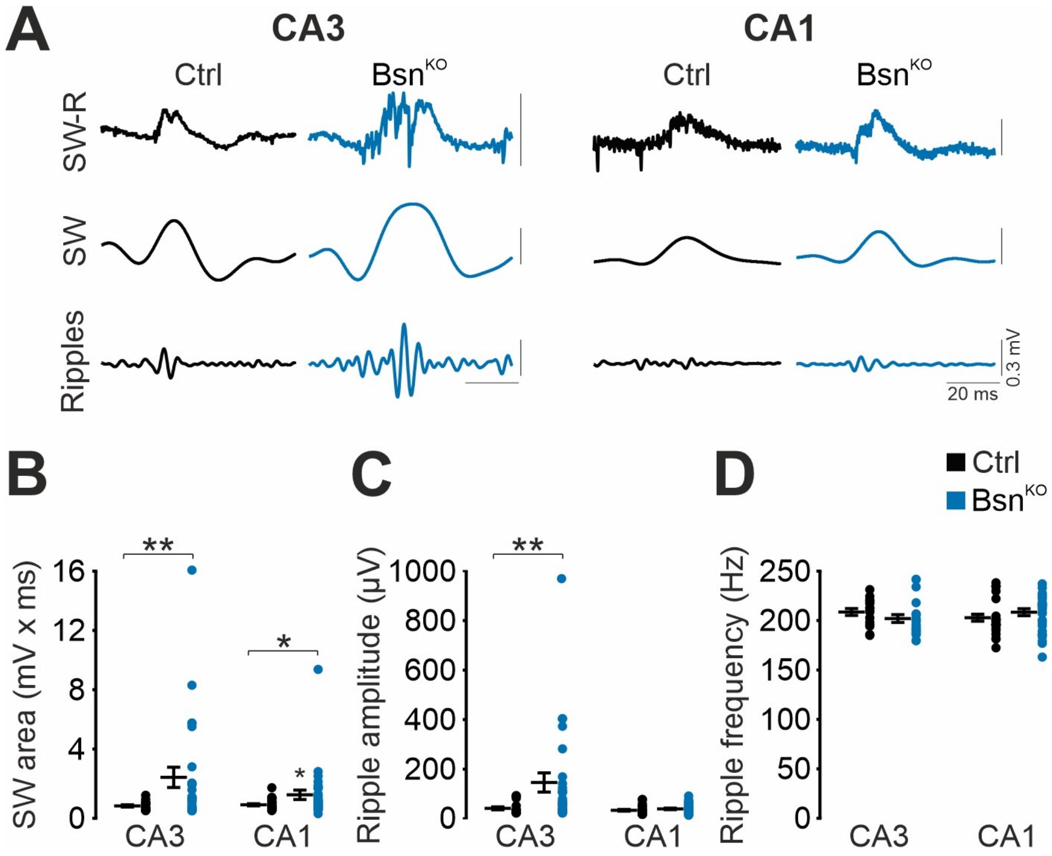


**Figure S3 (supplement to Figure 3).** Augmented sharp wave and ripples (SW-R) in the hippocampal CA3 region of *Bsn*$^{KO}$ mice. **A)** Representative SW-R traces recorded in the CA3 (Ctrl (= *Bsn*$^{lx/lx}$): *N* = 4 mice, *n* = 16 slices; *Bsn*$^{KO}$: *N* = 5 mice, *n* = 27 slices) and CA1 (Ctrl: *N* = 5 mice, *n* = 26 slices; *Bsn*$^{KO}$: *N* = 5 mice, *n* = 33 slices) from Ctrl and *Bsn*$^{KO}$ mice with low-pass filtered (middle trace; <45 Hz) SW component and band-pass filtered (bottom trace; 120-300 Hz) ripple component. Summary graphs showing increased **B)** SW areas and **C)** ripple amplitudes and no alteration in the **D)** ripple frequencies in the CA3 and CA1 subregions of *Bsn*$^{KO}$ mice. Data presented as mean ± SEM. **B, C:** Mann-Whitney U test; **D: CA3:** Mann-Whitney U test; **CA1:** Student´s two-tailed test. *$P$ < 0.05, **$P$ < 0.01.

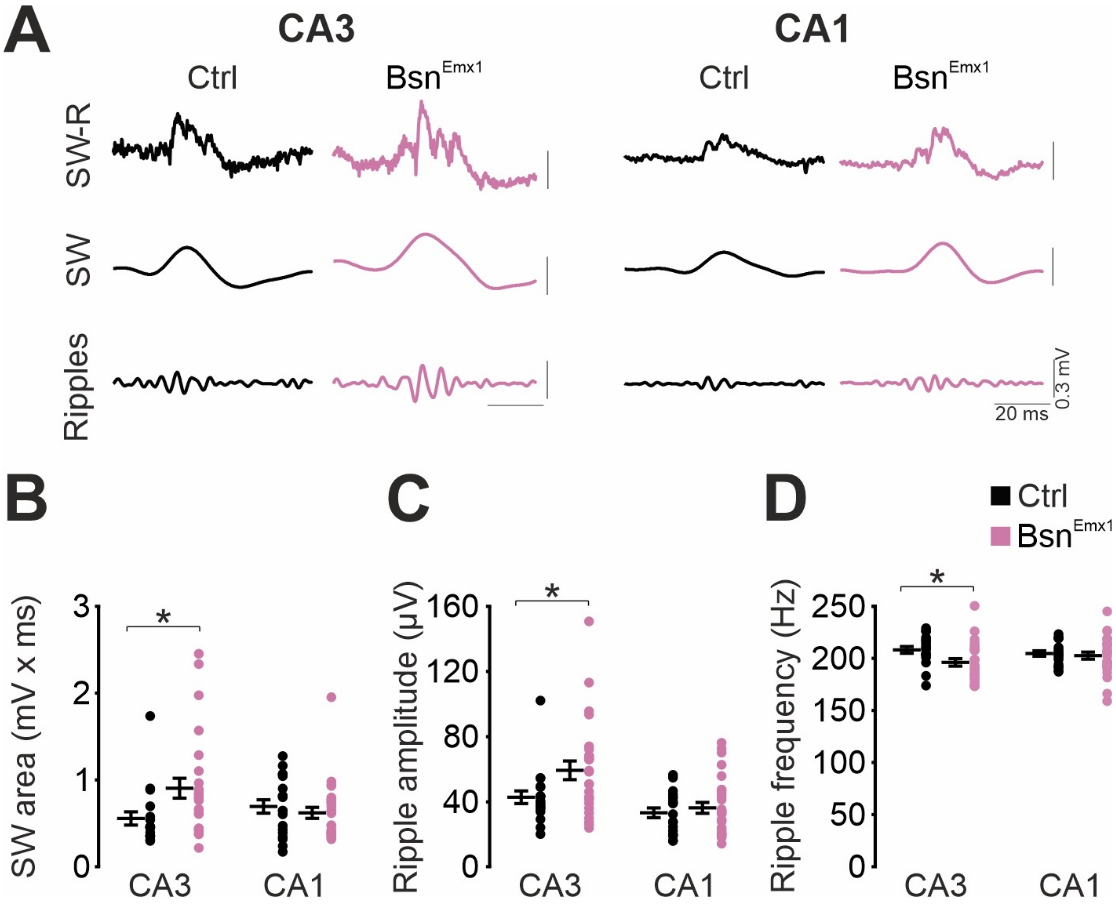


**Figure S4 (supplement to Figure 3).** Augmented sharp wave and ripples (SW-R) in the hippocampal CA3 region of *Bsn*$^{Emx1}$ mice. **A)** Representative SW-R traces recorded in the CA3 (Ctrl (= *Bsn*$^{lx/lx}$): *N* = 5 mice, *n* = 19 slices; *Bsn*$^{Emx1}$: *N* = 5 mice, *n* = 26 slices) and CA1 (Ctrl: *N* = 5 mice, *n* = 19 slices; *Bsn*$^{Emx1}$: *N* = 5 mice, *n* = 28 slices) from Ctrl and *Bsn*$^{Emx1}$ mice with low-pass filtered (middle trace; <45 Hz) SW component and band-pass filtered (bottom trace; 120-300 Hz) ripple component. Summary graphs showing increased **B)** SW areas and **C)** ripple amplitudes and a reduction in the **D)** ripple frequencies in the CA3 subregion of *Bsn*$^{Emx1}$ mice. Data presented as mean ± SEM. **B: CA3:** Student´s two-tailed test**, CA1:** Mann-Whitney U test; **C:** Mann-Whitney U test; **D:** Student´s two-tailed test. *$P$ < 0.05.

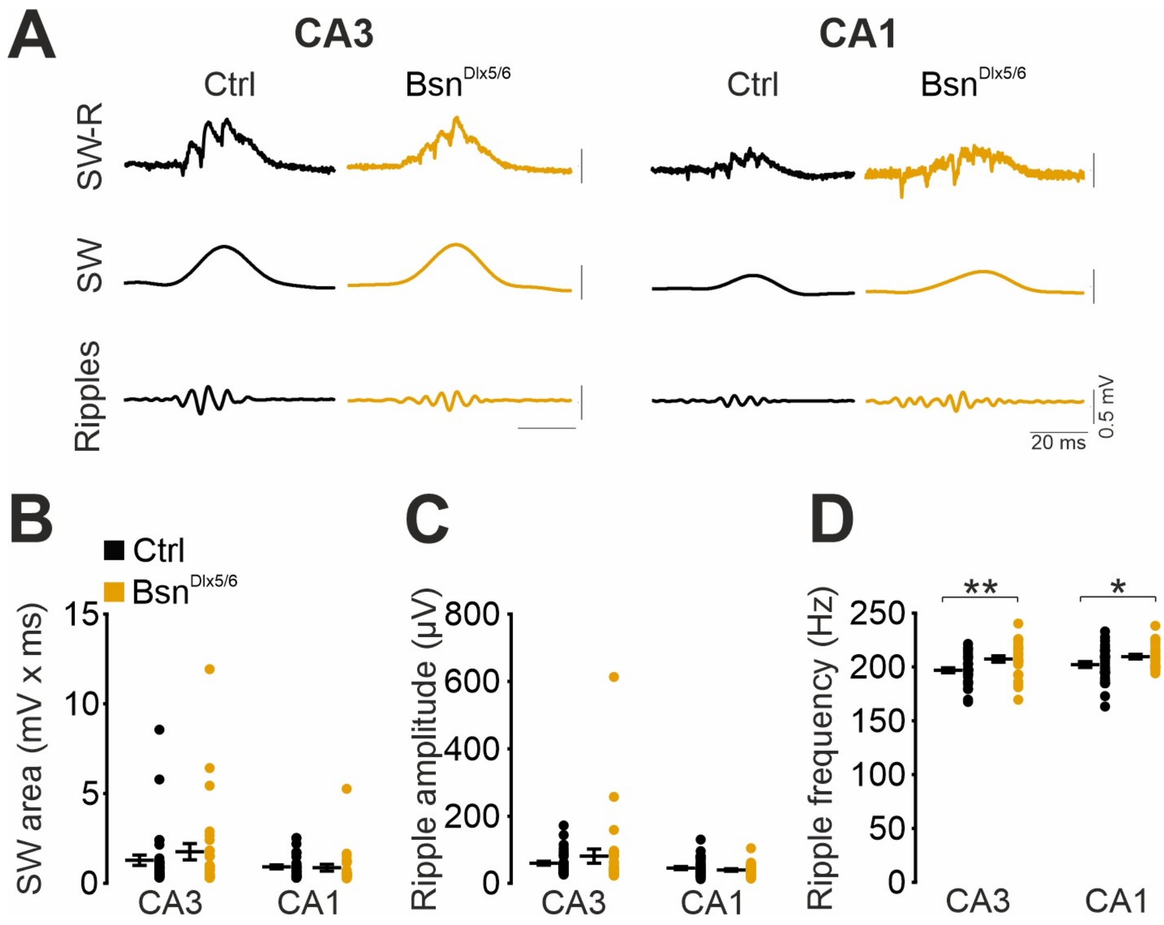


**Figure S5 (supplement to Figure 3).** Increased ripple frequency in the hippocampal CA3 / CA1 regions of $Bsn^{Dlx5/6}$ mice. **A)** Representative sharp wave and ripple (SW-R) traces recorded in the CA3 (Ctrl (= $Bsn^{lx/lx}$): *N* = 8 mice, *n* = 33 slices; cKO: *N* = 8 mice, *n* = 29 slices) and CA1 (Ctrl: *N* = 8 mice, *n* = 34 slices; cKO: *N* = 8 mice, *n* = 27 slices) from Ctrl and $Bsn^{Dlx5/6}$ mice with low-pass filtered (middle trace; <45 Hz) SW component and band-pass filtered (bottom trace; 120-300 Hz) ripple component. Summary graphs showing no statistical change in the **B)** SW areas and **C)** ripple amplitudes but an increase in the **D)** ripple frequencies in the CA3 and CA1 subregions of $Bsn^{Dlx5/6}$ mice. Data presented as mean ± SEM. **B, C**: Mann-Whitney U test; **D**: Student´s two-tailed test. *$P < 0.05$, **$P < 0.01$.

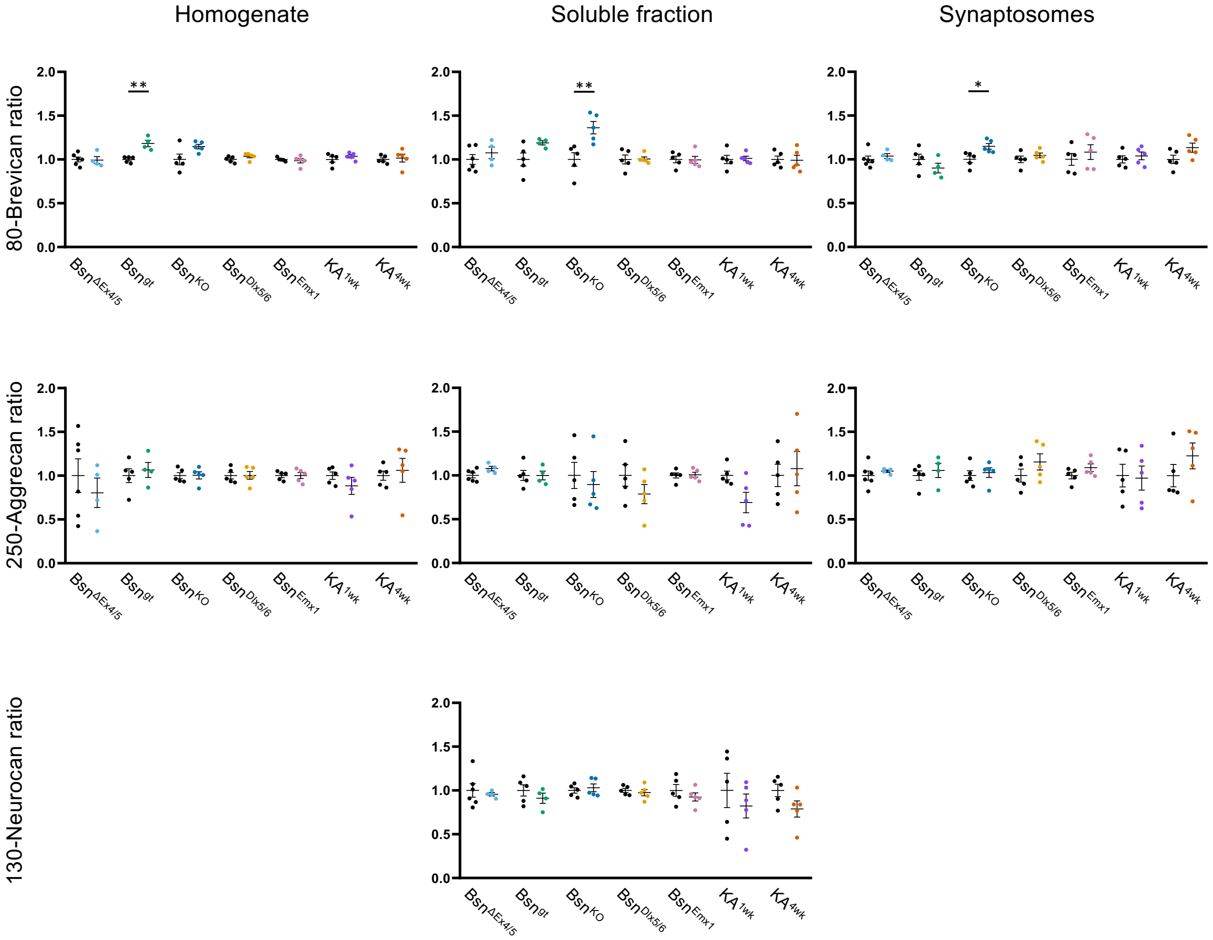


**Figure S6 (supplement to Figure 4).** Normalized ratios of cleaved to total proteoglycans in subcellular forebrain homogenate, soluble and synaptosomal fractions from the epilepsy models. Data points indicate the normalized ratios of 80kDa C-terminal Brevican fragment to total Brevican (upper row), of 250kDa N-terminal Aggrecan fragment to total Aggrecan, and of 130kDa N-terminal neurocan fragment to total neurocan. Statistics: Two-way ANOVA, Sidak‘s multiple comparisons test of treated vs. ctrl. $**P < 0.01$, $***P < 0.001$.

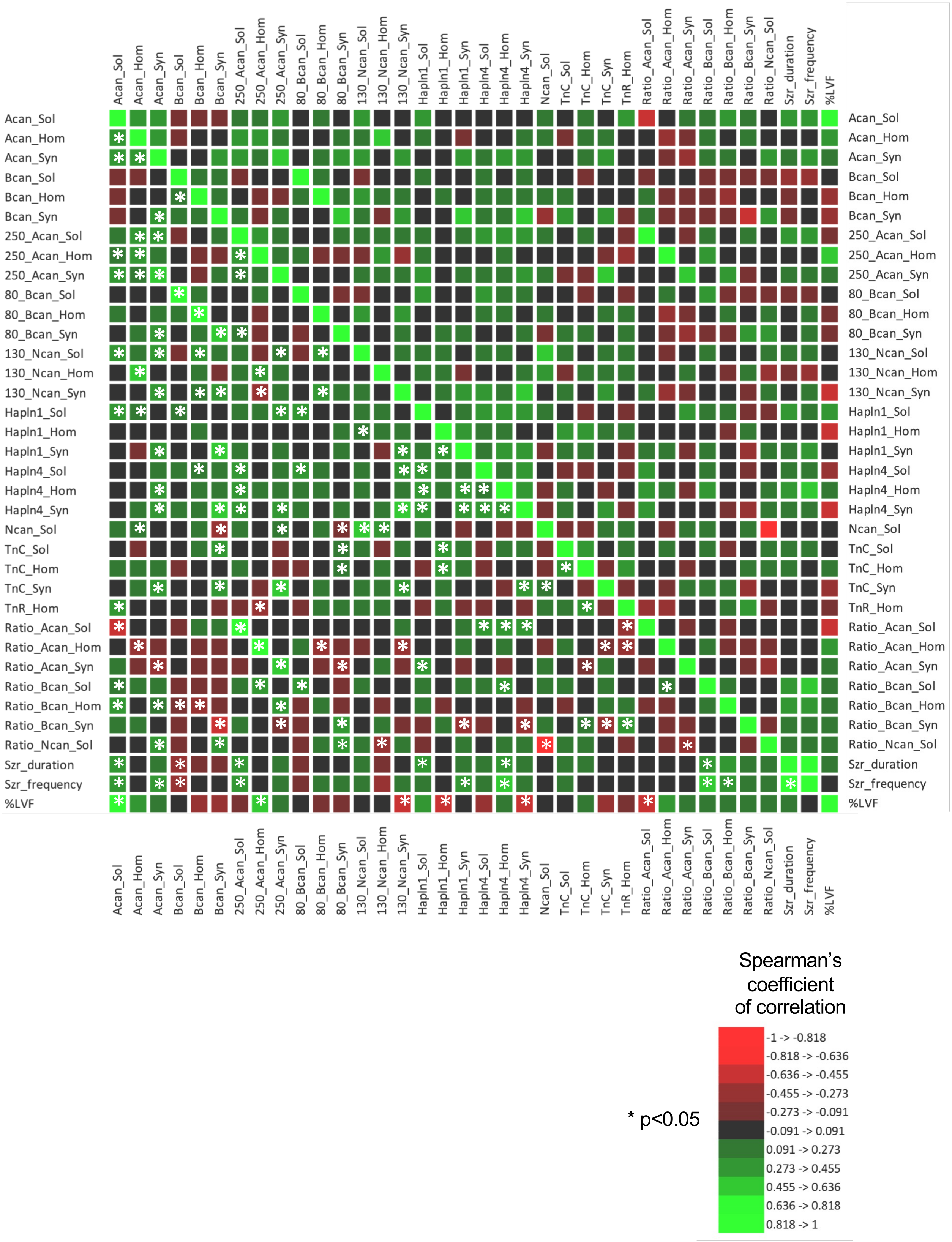


**Fig. S7 (supplement to Figures 4, 5)** Color-coded values of Spearman coefficients of correlation between all ECM and seizure parameters. To facilitate color perception, * $P < 0.05$ are shown only for one half of the correlation matrix, as it is symmetric.

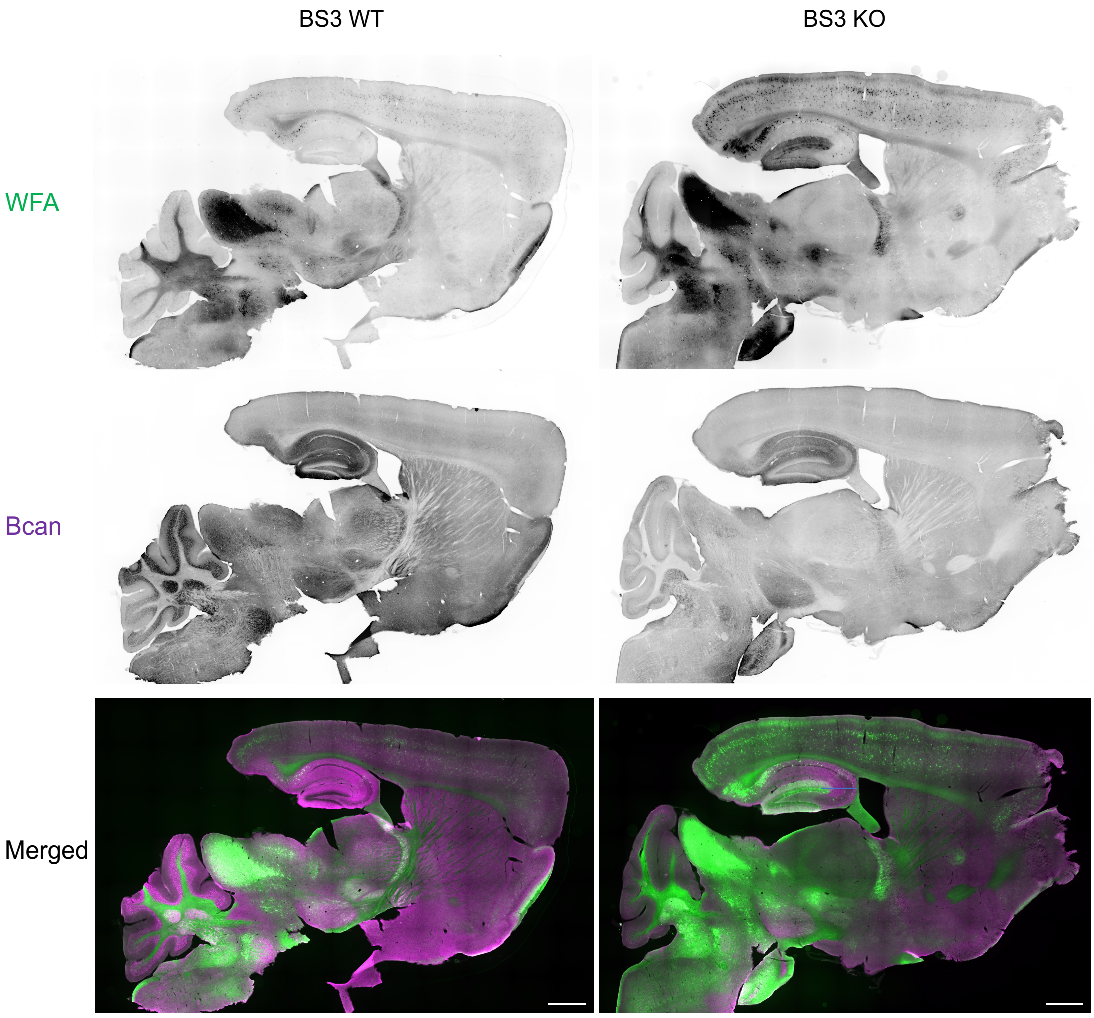


**Fig. S8 (supplement to Figure 6)** Lectin- and immuno-histochemical distribution of Wisteria floribunda agglutinin (WFA; green) and Brevican (Bcan; purple) in saggital brain sections of a *Bsn*$^{KO}$ mouse. A floxed mouse *Bsn*$^{lx/lx}$ served as control. Scale bar: 1 mm. Medial/Lateral (ML): 1,56 mm.

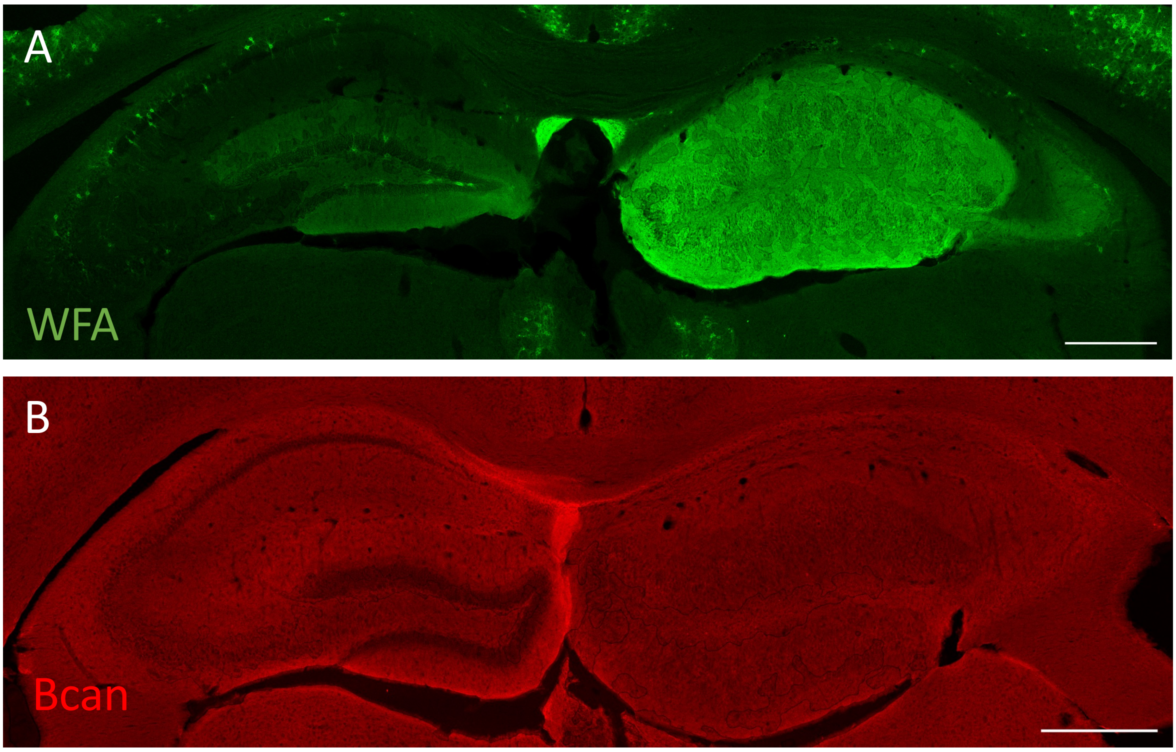


**Figure S9 (supplement to Figure 6).** Distribution of (**A**) *Wisteria floribunda* agglutinin (WFA) binding sites (green) and (**B**) Brevican (Bcan) immunoreactivity (red) in frontal brain sections of $KA^{4wk}$ mice. The right side was injected, the left side served as control. WFA binding is massively increased in the hippocampus 4 weeks after kainate injection (**A**) while Brevican levels are reduced (**B**). Scale bar: 1000 μm.
